# Fiber Microstructure Quantile (FMQ) Regression: A Novel Statistical Approach for Analyzing White Matter Bundles from Periphery to Core

**DOI:** 10.1101/2024.10.19.619237

**Authors:** Zhou Lan, Yuqian Chen, Jarrett Rushmore, Leo Zekelman, Nikos Makris, Yogesh Rathi, Alexandra J. Golby, Fan Zhang, Lauren J. O’Donnell

## Abstract

The structural connections of the brain’s white matter are critical for brain function. Diffusion MRI tractography enables the in-vivo reconstruction of white matter fiber bundles and the study of their relationship to covariates of interest, such as neurobehavioral or clinical factors. In this work, we introduce Fiber Microstructure Quantile (FMQ) Regression, a new statistical approach for studying the association between white matter fiber bundles and scalar factors (e.g., cognitive scores). Our approach analyzes tissue microstructure measures based on *quantile-specific bundle regions*. These regions are defined in a data-driven fashion according to the quantiles of fractional anisotropy (FA) of a *population fiber bundle,* which pools all individuals’ bundles. The FA quantiles induce a natural subdivision of a fiber bundle, defining regions from the periphery (low FA) to the core (high FA) of the population fiber bundle. To investigate how fiber bundle tissue microstructure relates to covariates of interest, we employ the statistical technique of quantile regression. Unlike ordinary regression, which only models a conditional mean, quantile regression models the conditional quantiles of a response variable. This enables the proposed analysis, where a quantile regression is fitted for each quantile-specific bundle region. To demonstrate FMQ Regression, we perform an illustrative study in a large healthy young adult tractography dataset derived from the Human Connectome Project-Young Adult (HCP-YA), focusing on particular bundles expected to relate to particular aspects of cognition and motor function. In comparison with traditional regression analyses based on FA Mean and Automated Fiber Quantification (AFQ), we find that FMQ Regression provides a superior model fit with the lowest mean squared error. This demonstrates that FMQ Regression captures the relationship between scalar factors and white matter microstructure more effectively than the compared approaches. Our results suggest that FMQ Regression, which enables FA analysis in data-driven regions defined by FA quantiles, is more powerful for detecting brain-behavior associations than AFQ, which enables FA analysis in regions defined along the trajectory of a bundle. FMQ Regression finds significant brain-behavior associations in multiple bundles, including findings unique to males or to females. In both males and females, language performance is significantly associated with FA in the left arcuate fasciculus, with stronger associations in the bundle’s periphery. In males only, memory performance is significantly associated with FA in the left uncinate fasciculus, particularly in intermediate regions of the bundle. In females only, motor performance is significantly associated with FA in the left and right corticospinal tracts, with a slightly lower relationship at the bundle periphery and a slightly higher relationship toward the bundle core. No significant relationships are found between executive function and cingulum bundle FA. Our study demonstrates that FMQ Regression is a powerful statistical approach that can provide insight into associations from bundle periphery to bundle core. Our results also identify several brain-behavior relationships unique to males or to females, highlighting the importance of considering sex differences in future research.

## 1. Introduction

The white matter plays a critical role in brain function, serving as the brain’s communication infrastructure that is essential for the proper functioning of various cognitive domains (Fields, 2008). Diffusion magnetic resonance imaging (dMRI) is an advanced imaging technique that can measure the diffusion process of water molecules and facilitate the investigation of white matter. dMRI tractography is a three-dimensional reconstruction technique to reconstruct white matter fiber bundles using data collected by dMRI (Basser et al., 2000). Many large white matter fiber bundles have a long history of anatomical study and are classically defined (e.g., the arcuate fasciculus and the corticospinal tract). Recent machine learning methods can use dMRI tractography to efficiently identify white matter fiber bundles of individuals (Garyfallidis et al., 2012; Wasserthal et al., 2018; F. Zhang et al., 2018). The fiber bundles obtained from tractography enable the quantitative study of the brain’s white matter anatomy (F. Zhang et al., 2022) and its associations with scalar factors, such as those describing individual cognition or behavior (e.g., language, memory, executive function, or motor) (Zekelman et al., 2022), or those describing diseases or disorders (Damatac et al., 2022; Kruper et al., 2023).

Analyzing the association between fiber bundles and scalar factors requires summary data derived from fiber bundles. One popular quantity for fiber bundle analysis is fractional anisotropy (FA), a scalar value between zero and one that describes the degree of anisotropy of a diffusion process and relates to the geometry and health of the tissue (Basser & Pierpaoli, 2011). The FA mean within the fiber bundle has been widely used due to its parsimony (Bozzali et al., 2002; Ciccarelli et al., 2003; Schilling et al., 2023; Zekelman et al., 2022; F. Zhang et al., 2022). More sophisticated summary data can provide profiles that describe data along fiber bundles (Batchelor et al., 2006; Chandio et al., 2020; Colby et al., 2012; Corouge et al., 2004, 2006; Gerig et al., 2004; L. J. O’Donnell et al., 2009; Yeatman et al., 2012). For example, the Automated Fiber Quantification (AFQ) method produces an FA profile along a fiber bundle (Yeatman et al., 2012). This popular method has enabled clinical research applications (Johnson et al., 2022; Kruper et al., 2023, 2024; Sarica et al., 2017; Schilling, Archer, et al., 2022) and is undergoing active research (Gari et al., 2023; Neher et al., 2023).

The above summary data has limitations in analyzing the associations between fiber bundles and scalar factors. The FA mean overlooks the known microstructural variations of FA within a fiber bundle due to factors such as crossing or fanning white matter geometry and axons entering and leaving the bundle (Colby et al., 2012; Jeurissen et al., 2013; Jones, 2010; Jones et al., 2013; L. J. O’Donnell et al., 2009; Schilling, Tax, et al., 2022). Methods such as AFQ analyze FA at each cross-sectional location along the bundle profile (Chandio et al., 2020; Yeatman et al., 2012), which causes challenges in capturing the microstructural variations in the cross-section of the bundle, such as those due to axons crossing, entering, or leaving the bundle. It is also challenging to align bundle profiles across subjects in the presence of anatomical variability in bundle size and shape (Chamberland et al., 2019; Maddah et al., 2006).

In our paper, we are motivated to address the aforementioned limitations. Our methodology is built on a *population fiber bundle* (Chenot et al., 2019; Elias et al., 2024; L. O’Donnell & Westin, 2005; F. Zhang et al., 2018) that pools all individuals’ fiber bundles. We use the quantiles of FA within the population fiber bundle to define regions that we call *quantile-specific bundle regions*. As can be observed in Figure 1, the quantile-specific bundle regions finely subdivide the bundle and generally range from the central portion of a fiber bundle with higher FA (i.e., the bundle core) to more peripherally located regions of a fiber bundle with lower FA (i.e., the bundle periphery). In contrast to other methods for studying fiber tract microstructure, which represent fiber bundles using spatial or geometric information (such as subdivisions along bundle length or medial models of the bundle core (Yushkevich et al., 2009), our quantile-specific bundle regions are a data-driven approach that directly studies the microstructure within the tract. Here, we observe that the data-driven quantiles induce a natural subdivision of a fiber bundle according to the spatial distribution of its tissue microstructure.

**Figure 1:**
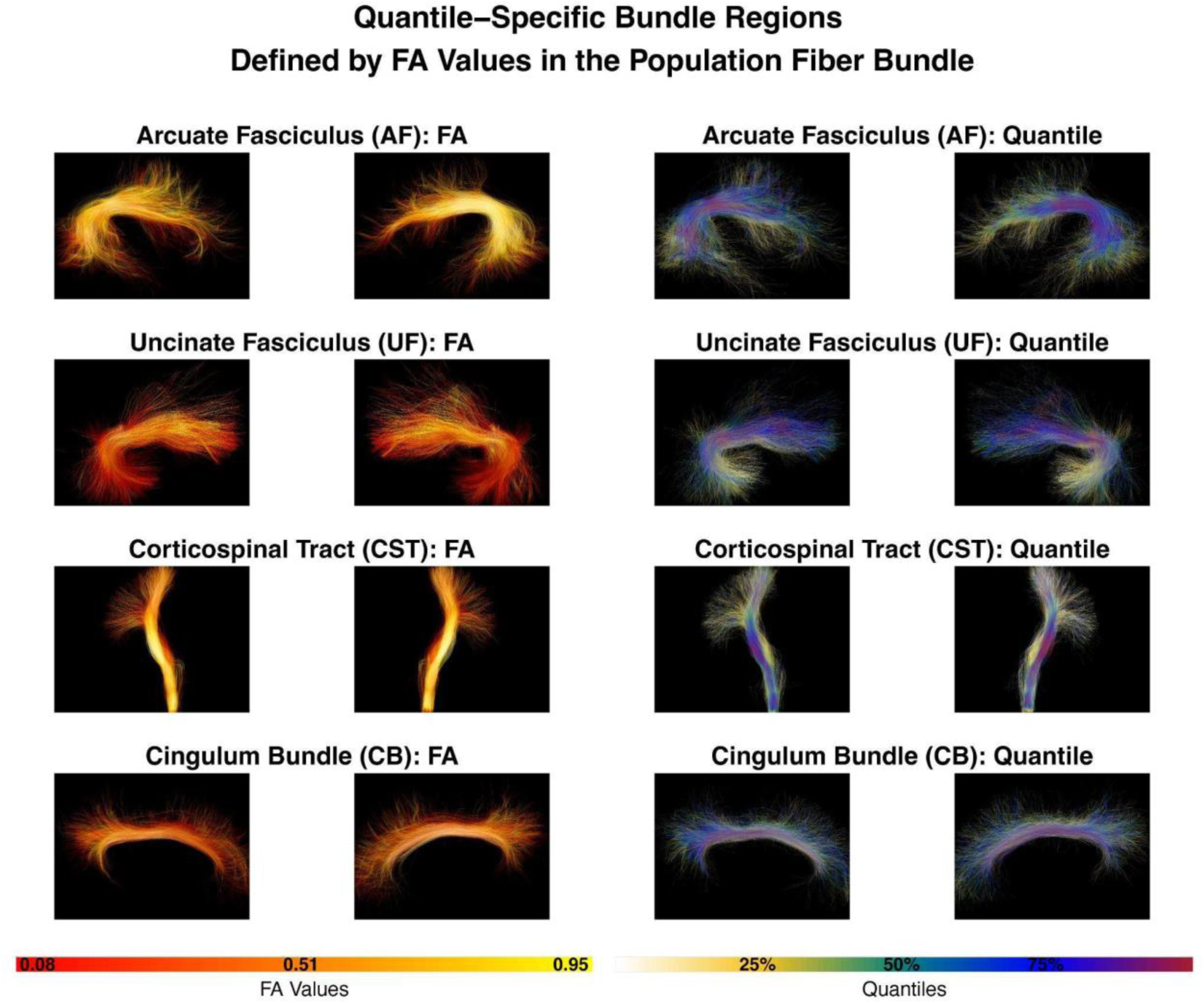
The *population fiber bundles* studied in this paper are the Arcuate Fasciculus (AF), Uncinate Fasciculus (UF), Corticospinal Tract (CST), and Cingulum Bundle (CB), shown here in both the left and right hemispheres. In each row, the two left images show fiber tracts (right and left hemispheres) colored by FA values, and the two right images show the corresponding quantiles on the fiber tracts. In this way, the FA values define *quantile-specific bundle regions*. At the bottom of the figure, we provide the color bars for FA values and quantiles.

In this paper, we employ quantile regression, a popular statistical technique, to enable the investigation of FA in the proposed quantile-specific bundle regions, providing insights into the association of microstructure with scalar factors. Quantile regression is a robust, semi-parametric statistical approach that models conditional quantiles of response variables (Koenker et al., 2020) and does not make strict assumptions about the error distribution (Wei et al., 2006). In contrast to ordinary regression, which models the association between the conditional mean of the response variable and the independent variables, quantile regression allows the investigation of associations between quantiles of the response variable and the independent variables.

In our proposed approach, we consider that FA within the population fiber bundle is the response variable, and the scalar factors describing neurobehavioral function are the independent variables. We employ quantile regression to study the conditional quantiles of FA within the population fiber bundle and their associations with the scalar factors. We call our proposed approach *Fiber Microstructure Quantile (FMQ) Regression*. While quantile regression has been successfully applied in many fields, including genetics and environmental science (Koenker et al., 2020), the proposed FMQ Regression is the first paper to use quantile regression to perform microstructural white matter analysis. To demonstrate its performance, we provide an illustrative study investigating sex-specific effects in brain-behavior associations using a large dataset.

In the rest of the paper, we first introduce the dataset for our illustrative study. We further present our proposed method in Section 2. Section 3 provides the results of our illustrative study using our methodology and other methods. Section 4 discusses the results of our illustrative study and the differences between methods and provides conclusions.

## 2. Methods

In this section, we first describe the tractography dataset used to illustrate the proposed method (Section 2.1). Then, in Section 2.2, we describe the FMQ Regression method that employs fiber tract data and quantile regression for population-based inference. Next, in Section 2.3, we describe two popular current methods that are used for comparison. Finally, in Section 2.4, we describe the statistical estimates resulting from the three compared methods and the approach for results visualization.

### 2.1 HCP-YA Dataset for Illustrative Study

To demonstrate our proposed approach, we perform an illustrative study based on a large tractography dataset, focusing on specific tracts expected to relate to particular aspects of motor function and cognition as described in a recent review (Forkel et al., 2022). We use dMRI and scalar factors (i.e., neurobehavioral assessments of language, memory, executive function, and motor performance) from the Human Connectome Project-Young Adult (HCP-YA), a comprehensive multimodal dataset acquired from healthy young adults (Van Essen et al., 2013) that provides minimally processed dMRI (Glasser et al., 2013). HCP-YA is publicly available and de-identified in accordance with the Health Insurance Portability and Accountability Act (HIPAA) Privacy Rules. The data were accessed and used in compliance with the HCP Data Use Terms (https://www.humanconnectome.org/study/hcp-young-adult/data-use-terms), which prohibit attempts to re-identify participants and require responsible data stewardship. All analyses were conducted in accordance with these terms, and no efforts were made to identify individual participants.

We use tractography data previously computed for a cohort of 809 HCP-YA participants published in (Zekelman et al., 2022). The study dataset comprises 809 participants, with 382 males and 427 females. Their ages range from 22 to 36 years, with an average age of 28.6. Brief details about data acquisition and processing follow. The HCP-YA dataset (Glasser et al., 2013) was acquired using three shells (b = 1000, 2000, and 3000 s/mm²), with TE/TR = 89.5/5520 ms and an isotropic voxel size of 1.25 mm³. The b = 3000 shell, consisting of 90 gradient directions and all b = 0 scans, was extracted to reduce computation time while providing high angular resolution for tractography (F. Zhang et al., 2018). Whole brain tractography was computed by applying a two-tensor Unscented Kalman Filter (UKF) method (Reddy & Rathi, 2016), which is effective at reconstructing white matter tracts across various dMRI acquisitions and the lifespan (F. Zhang et al., 2018) with advantages for reconstructing anatomical somatotopy (He et al., 2023). UKF tractography used a two-tensor model to account for crossing fibers (Farquharson et al., 2013; Vos et al., 2013) and provided fiber-bundle-specific microstructural measures from the first tensor, which modeled the tract being traced (Reddy & Rathi, 2016). White matter tracts were identified for each subject using the white matter analysis machine learning approach that can robustly identify white matter tracts across the human lifespan, health conditions including brain tumors, and different image acquisitions (Cetin-Karayumak et al., 2024; F. Zhang et al., 2018) with high test-retest reproducibility (F. Zhang et al., 2019).

In our illustrative study, we investigate the arcuate fasciculus (AF), uncinate fasciculus (UF), cingulum (CB), and corticospinal tract (CST). These four white matter tracts of interest are investigated in each subject’s left and right hemispheres. Each fiber tract contains a collection of streamlines representing the pathway of a particular white matter connection. Each streamline is composed of a sequence of points (*streamline points*) and their associated FA values. In this paper, we use FA as a primary measure for tract analysis, though the methods we propose are equally applicable to other microstructure or imaging data measured within fiber tracts, e.g., mean diffusivity (MD) (Basser, 1995) or neurite orientation dispersion and density imaging (NODDI) (Daducci et al., 2015; H. Zhang et al., 2012). (In the supplementary materials, we also provide the illustrative study based on MD.)

We assess the relationship between the FA of each fiber tract and selected scalar factors, as summarized in Table 1. We study the associations of the AF, UF, CB, and CST with scalar factors of language, memory, executive function, and motor, respectively. Our choices of fiber tracts and corresponding scalar factors follow a recent review of fiber tracts and potentially associated neuro-behavioral functions in health and disease based on the existing literature (Forkel et al., 2022). In our work, the scalar factors are assessments from the NIH Toolbox, the state-of-the-art for neurobehavioral measurement (Hodes et al., 2013). These include the NIH Toolbox Oral Reading Recognition Test (Gershon et al., 2014), Picture Vocabulary Test (Gershon et al., 2014), Picture Sequence Memory Test (Loring et al., 2019), List Sorting Working Memory Test (Tulsky et al., 2014), Dimensional Change Card Sort Test (Zelazo et al., 2014), Flanker Inhibitory Control and Attention Test (Zelazo et al., 2014), 2-minute Walk Endurance Test (Reuben et al., 2013), and 4-Meter Walk Gait Speed Test (Reuben et al., 2013).

**Table 1:** Input data for the illustrative study includes fiber tracts, corresponding neurobehavioral functions following a recent review, and scalar factors from the NIH Toolbox. We include an abbreviated name for each scalar factor.

| Fiber Tract | Neuro-behavioral Function | Scalar Factor |
| --- | --- | --- |
| Arcuate Fasciculus (AF) | Language | Picture Vocabulary Test (PicVocab) |
|  |  | Oral Reading Recognition Test (ReadEng) |
| Uncinate Fasciculus<br>(UF) | Memory | Picture Sequence Memory Test (PicSort) |
|  |  | List Sorting Working Memory Test (ListSort) |
| Corticospinal Tract<br>(CST) | Motor | Walk Endurance Test (Endurance) |
|  |  | 4-Meter Walk Gait Speed Test (GaitSpeed) |
| Cingulum Bundle<br>(CB) | Executive<br>function | Dimensional Change Card Sort Test (CardSort) |
|  |  | Flanker Inhibitory Control and Attention Test (Flanker) |

### 2.2. Key Steps in FMQ Regression

#### 2.2.1 Step 1: Population Fiber Bundle Construction

Population fiber bundle construction is the fundamental step that allows us to perform quantile-based microstructural analysis. To construct a population fiber bundle, the bundle and its FA values must first be identified in all subjects in the population. This can be achieved using methods such as virtual dissection (Catani et al., 2002) or automatic segmentation (Garyfallidis et al., 2018; Vázquez et al., 2020; F. Zhang et al., 2018). The resulting individual fiber bundles have different numbers of streamlines, and their shapes and lengths are not the same due to factors such as anatomical variability and neural plasticity. Constructing a population fiber bundle, which contains the amalgamated streamlines among all individuals within a cohort (Chenot et al., 2019; Elias et al., 2024; L. J. O’Donnell et al., 2012; L. O’Donnell & Westin, 2005), is the first step in microstructural inference using FMQ. Population fiber tracts studied in this paper are given in Figure 1. The statistical inference and the following anatomical interpretations rely on the constructed population fiber bundle.

We use the notation *i* ∈ {1,2, …, *I*} to denote an individual, where *I* is the total number of individuals. For a certain fiber tract, we randomly select *K* streamlines from each individual’s fiber tract (i.e., sampled fiber tract) to ensure each individual’s fiber tract has equal “weight” in constructing the population fiber bundle. We use 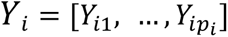 to denote the microstructure measures (FA values) of each individual, where *p*_*i*_ is the number of streamline points of the *i*-th individual’s sampled fiber tract. We construct the population fiber bundle by *assembling* individuals’ fiber tracts *together*. The assembled data thus is a population fiber bundle with FA values ([*Y*_1_, …, *Y*_*I*_]). Figure 2 illustrates the process of population fiber bundle construction. This step makes our analysis very different from other quantile regression techniques that employ the FA mean of the entire fiber tract (Lv et al., 2021; Ryan et al., 2022).

**Figure 2:**
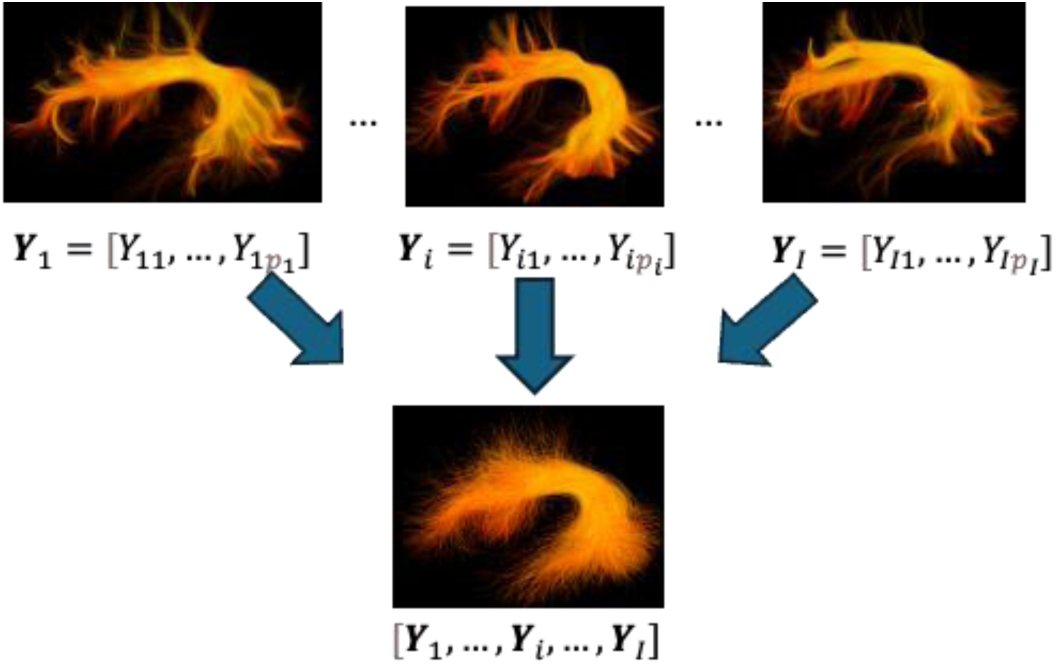
The top images show individual subject fiber tracts. By *assembling* individuals’ fiber tracts *together*, we create a population fiber bundle (bottom image) with FA values ([*Y*_1_, …, *Y*_*I*_]).

In our HCP-YA illustrative study, the total number of individuals is *I* = 809. For each individual, we randomly select *K* = 2000 streamlines (Wasserthal et al., 2019) for each individual’s fiber bundle. Therefore, *p*_*i*_ is determined by the individual’s sampled fiber tract.

#### 2.2.2 Step 2: Quantile-Specific Bundle Region Creation

In this step, we provide a data-driven approach to create quantile-specific bundle regions defined by the FA values in the population fiber bundle. Let *G*_*Y*_(*τ*) be the *τ*-th quantile of the population fiber bundle’s FA values ([*Y*_1_, …, *Y*_*I*_]). By using the values of *G*_*Y*_(*c*/*C*) for *c* ∈ {1, …, *C* − 1}, as cut-off values (the dashed lines in Figure 2), we can create *C quantile-specific bundle regions* ranging from *bundle periphery* to *bundle core* (Figure 2), and each quantile-specific bundle region has the same number of streamline points. We define 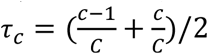 for *c* ∈ {1, …, *C* − 1}. For each quantile-specific bundle region *c* ∈ {1, …, *C*}, the value *G*_*Y*_(*τ*_*c*_) is defined as the *typical FA value* since it is the middle value between the boundary cut-offs (see blue arrows in Figure 3). The typical FA value indicates the representative FA value in the quantile-specific bundle region and is used as the dependent variable in the regression model introduced in Section 2.2.3.

**Figure 3:**
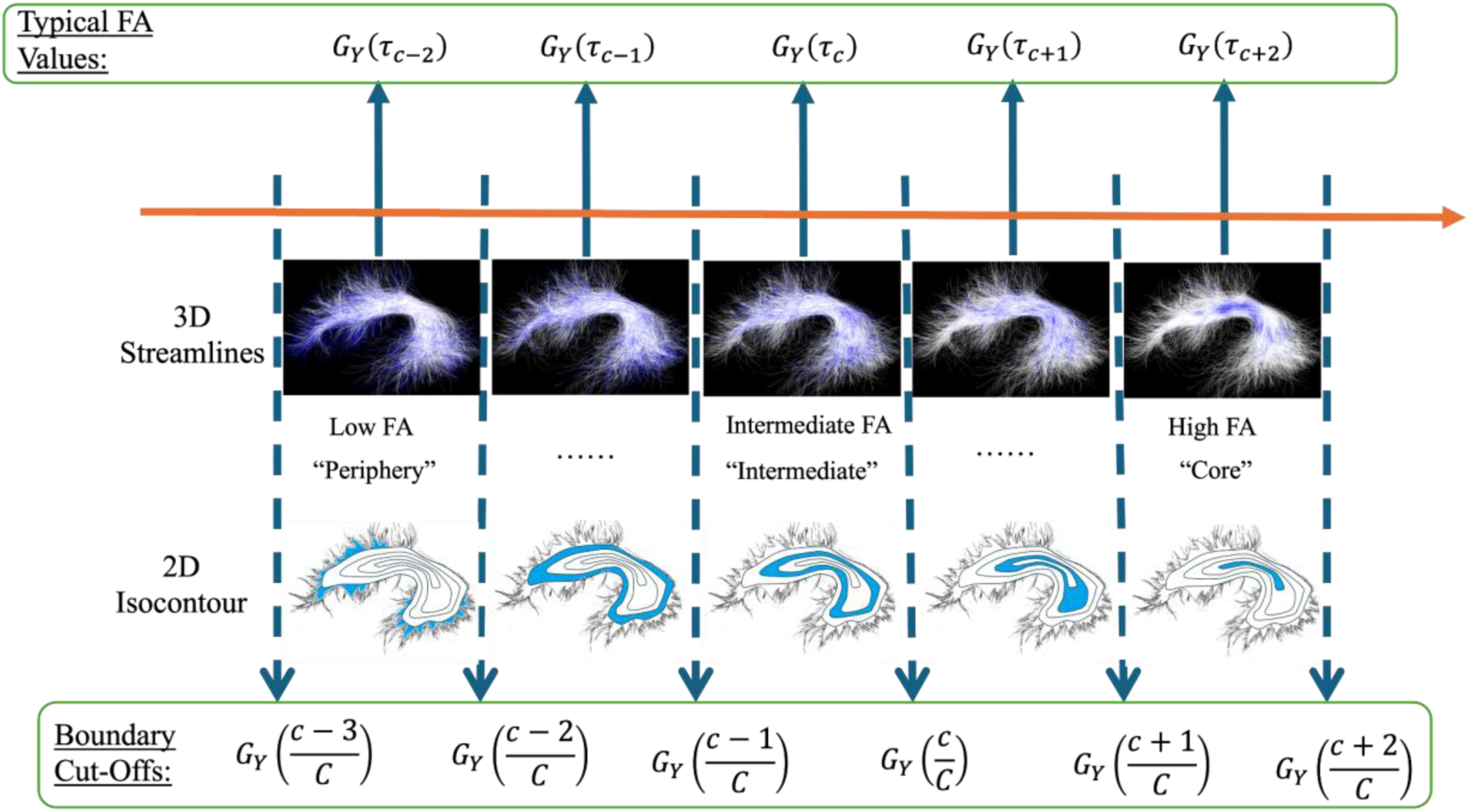
Graphical illustration of the population FA distribution sorted along the orange arrow from low to high FA (e.g., bundle periphery, to intermediate bundle regions, to bundle core). *C* quantile-specific bundle regions are defined based on the values of *G*_*Y*_(*c*/*C*) for *c* ∈ {1, …, *C* − 1}, as boundary cut-off values (the dashed lines). The upper and lower cut-off values give the range of FA within each quantile-specific bundle region. The value of *G*_*Y*_(*τ*_*c*_) is defined as the *typical FA value* in each region since it is the middle value between the boundary cut-offs. AF left is used as an illustrative example. Quantile-specific bundle regions are shown in blue in the 3D streamline data view (top) and in a schematic 2D isocontour diagram (bottom).

In our HCP-YA illustrative study, we choose *C* = 100, and thus the studied quantiles range from 0.5% to 99.5%. This choice of 100 corresponds to the default number of subdivisions in the AFQ method and is chosen for fair comparison (Yeatman et al., 2012).

#### 2.2.3 Step 3: Statistical Inference using Quantile Regression

In this section, we describe how to quantify the association between a typical FA value in a quantile-specific bundle region and a scalar factor. The relationship between the typical FA value and scalar factors is given as *G*_*Y*_(*τ*_*c*_) = *X*_*i*_*β*(*τ*_*c*_) where *X*_*i*_ is a covariate vector of an individual *i* containing the scalar factor. In the illustrative study of the HCP-YA data, we give the covariate vector of the regression model as 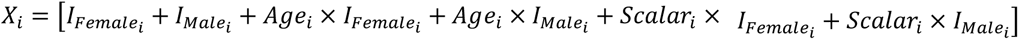. In the covariate vector, 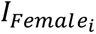 and 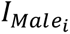 are indicator variables that equal 1 if the individual *i* is a female or a male, respectively. *Age*_*i*_ and *Scalar*_*i*_ are the values of the individual’s age and scalar factor, respectively. Without loss of generality, the scalar factor is scaled into the range between 0 and 1. This scaling step allows direct comparison of the magnitudes of the regression coefficients of different scalar factors. In this regression model, the intercepts and regression coefficients are computed for both males and females. The regression coefficient associated with the assessment, *β*(*τ*_*c*_), is used to quantify the effect of the scalar factor on the typical FA value of each quantile-specific bundle region.

The most conventional and fastest approach to estimating the coefficient vector is defined as follows,

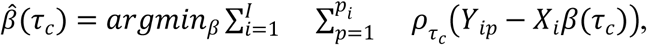

where 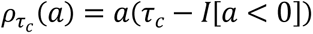 is the check function and *I*[*e*] is the indicator function for the event of *e*. By giving asymmetric weights to positive and negative values, the check function is a loss function to estimate *β*(*τ*_*c*_), making *X*_*i*_*β*(*τ*_*c*_) as close to the *τ*_*c*_-th quantile of [*Y*_1_, …, *Y*_*I*_]as possible. This estimator pools all FA values from all the individuals but does not consider the possible effect of between-individual variations. This estimator has been proven to be asymptotically consistent under certain conditions (Parente & Santos Silva, 2016):

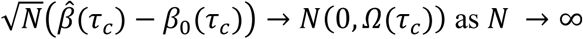

In other words, a large sample size of individuals (*N*) produces an estimator that better converges to the true regression coefficient values, i.e., *β*_0_(*τ*_*c*_). The asymptotic covariance matrix *Ω*(*τ*_*c*_) = *B*(*τ*_*c*_)^−1^*A*(*τ*_*c*_)*B*(*τ*_*c*_)^−1^allows us to make valid statistical inferences to account for between-individual variations, providing valid uncertainties of regression coefficient estimates. Furthermore, *Ω*(*τ*_*c*_) = *B*(*τ*_*c*_)^−1^*A*(*τ*_*c*_)*B*(*τ*_*c*_)^−1^ is feasible to be estimated using the data, denoted as 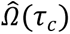 thus, we can avoid the computationally expensive bootstrap for obtaining the covariance of the estimate. The estimators for *A*(*τ*_*c*_) and *B*(*τ*_*c*_) can be found in Section 2.2 in Parente & Santos Silva, (2016). The Z-scores and adjusted p-values for the regression coefficient associated with a scalar factor are reported as the principal results. Benjamini–Hochberg-False Discovery Rate (BH-FDR) correction is applied to adjust p-values over the quantile-specific bundle regions for each scalar factor. We reject the null hypothesis if the corresponding p-value is smaller than 0.05. The Z-scores pool regression coefficient estimates’ magnitudes and uncertainties; thus they are used as the primary quantities to describe the effects of scalar factors.

### 2.3. Alternative Methods to be Compared

In this section, we provide alternative methods to be compared to our proposed method. We use the FA mean and AFQ tract profile as quantities or profiles measured from the fiber tract, and we build regression models based on these. The names of the two compared regression methods are FA Mean Regression (Section 2.3.1) and AFQ Regression (Section 2.3.2).

#### 2.3.1 FA Mean Regression

In FA Mean Regression, the fiber tract FA mean is the response of the regression models. The fiber tract FA mean is a value averaging all the FA values over streamline points within a fiber tract (L. J. O’Donnell & Westin, 2007; Zekelman et al., 2022). FA Mean Regression is the simplest method that uses the FA mean as an imaging biomarker. Simplicity makes it easy to use, but the detailed information on the microstructure within the tract is aggregated. The regression model is defined as *M*_*i*_ = *X*_*i*_*β* + *ε*_*i*_; *ε*_*i*_ ∼ *N*(0, *σ*^2^), where the tract FA mean is expressed as *M*_*i*_. In the regression, the covariate vector *X*_*i*_ is the same as in Section 2.2.3. The regression coefficient associated with each scalar factor quantifies the magnitude of the effect on the FA mean *M*_*i*_. The T-scores and p-values for the regression coefficient associated with a scalar factor are reported as the principal results. The p-values here are not corrected since there is no multiple comparison. We reject the null hypothesis if the corresponding p-value is smaller than 0.05. The T-scores that pool regression coefficient estimates’ magnitudes and uncertainties are used as the primary quantities to describe the effects of scalar factors.

#### 2.3.2 AFQ Regression

In AFQ Regression, FA values within the AFQ tract profile are the responses of regression models. The AFQ method (Yeatman et al., 2012) implemented in the dipy software (Garyfallidis et al., 2014) automatically computes the profile, which consists of mean FA values at sequential locations along a fiber tract. The AFQ tract profile of FA is individual-specific. While the fiber tracts are different between individuals, the *L* locations along a fiber tract can be matched across individuals, given their relative positions. We use *T*_*il*_ to denote the mean FA value of an individual *i* at a location *l*. We set up a linear regression as 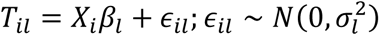. In the regression, the covariate vector *X*_*i*_ is the same as in Section 2.2.3. The regression coefficients associated with scalar factors quantify their effects on the FA at the location *l*. We set *L* = 100, which is the default convention for AFQ (Yeatman et al., 2012), following the tutorial instructions on the website^1^. This setting also makes it comparable to our FMQ Regression. The T-scores and adjusted p-values for the regression coefficient associated with a scalar factor are reported as the principal results. BH-FDR correction is applied to adjust p-values over the locations along a bundle for each scalar factor. We reject the null hypothesis if the corresponding p-value is smaller than 0.05. Similarly, the T-scores that pool regression coefficient estimates’ magnitudes and uncertainties are used as the primary quantities to describe the effects of scalar factors.

In this work, we perform both FMQ Regression and AFQ Regression using 100 regions. Though there may be some dependence between the hypothesis tests over these regions, the multiple comparison method we employ is reasonable. Specifically, a theoretical statistics study has proven that BH-FDR has good control of family-wise type I error even if the tests are dependent (Benjamini & Yekutieli, 2001).

### 2.4. Quantitative Comparison of Methods

We quantitatively measure the performance of FMQ Regression and other methods by comparing the mean squared error (MSE), a widely used metric in regression analysis due to its simplicity and ability to capture both the bias and variance of the estimators. The MSE of models measures the average squared difference between the estimated responses and the observed responses, providing a clear indication of the model’s predictive accuracy and fit to the data (Hastie et al., 2001). Lower MSE values signify models that closely align with the true underlying relationships, making the MSE a critical criterion for evaluating regression models. In FMQ Regression, the MSE was calculated for each quantile *τ*_*c*_ and defined as 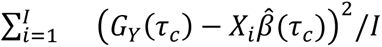. Similarly, in AFQ Regression, the MSE was calculated for each location *l* and defined as 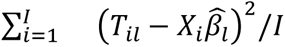. In FA Mean Regression, the MSE was defined as 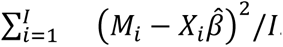.

## 3. Results

We first present the mean squared errors (MSE) results obtained from regression models assessing the associations between white matter tracts and scalar factors (Table 2). The MSE serves as a measure of the overall model fit, with lower values indicating better alignment between the model estimates and the observed data. Results show that FMQ Regression consistently produces the lowest MSE values compared to both FA Mean Regression and AFQ Regression. This finding suggests that the FMQ Regression method provides a superior model fit, capturing the relationship between the scalar factor and white matter tracts more effectively than the other approaches. Given this quantitative comparison result, which strongly favors the proposed FMQ Regression, we will next perform a further assessment of the regression results generated by the three methods in the illustrative study.

**Table 2.** MSE for FA Mean Regression, AFQ Regression, and FMQ Regression models. Each value represents the MSE, with standard deviations (over locations or quantiles) provided in parentheses for AFQ and FMQ Regression. In each row, the lowest MSE (best model fit) is shown in bold text.

| Fiber Bundle | Scalar Factor | FA Mean Regression | AFQ Regression | FMQ Regression |
| --- | --- | --- | --- | --- |
| AF Left | PicVocab | 4.90e-04 | 8.80e-04 (3.22e-04) | **1.34e-05 (8.04e-06)** |
|  | ReadEng | 4.88e-04 | 8.80e-04 (3.22e-04) | **1.19e-05 (6.14e-06)** |
| AF Right | PicVocab | 8.60e-04 | 8.35e-04 (3.08e-04) | **2.18e-05 (1.17e-05)** |
|  | ReadEng | 8.46e-04 | 8.37e-04 (3.10e-04) | **1.79e-05 (9.67e-06)** |
| UF Left | PicSEq | 1.08e-03 | 5.95e-04 (3.16e-04) | **2.21e-05 (1.65e-05)** |
|  | ListSort | 1.07e-03 | 5.94e-04 (3.17e-04) | **1.96e-05 (1.42e-05)** |
| UF Right | PicSEq | 1.17e-03 | 5.25e-04 (2.43e-04) | **2.30e-05 (1.88e-05)** |
|  | ListSort | 1.17e-03 | 5.25e-04 (2.44e-04) | **1.96e-05 (1.63e-05)** |
| CST Left | Endurance | 6.42e-04 | 4.81e-03 (2.13e-03) | **1.63e-05 (5.76e-06)** |
|  | GaitSpeed | 6.37e-04 | 4.83e-03 (2.14e-03) | **2.01e-05 (7.85e-06)** |
| CST Right | Endurance | 5.89e-04 | 4.51e-03 (1.99e-03) | **8.60e-06 (2.82e-06)** |
|  | GaitSpeed | 5.94e-04 | 4.51e-03 (1.99e-03) | **1.10e-05 (3.90e-06)** |
| CB Left | CardSort | 1.00e-03 | 1.87e-03 (1.77e-04) | **2.83e-05 (2.64e-05)** |
|  | Flanker | 9.95e-04 | 1.87e-03 (1.75e-04) | **2.82e-05 (2.65e-05)** |
| CB Right | CardSort | 1.06e-03 | 2.65e-03 (3.41e-04) | **8.90e-06 (7.19e-06)** |
|  | Flanker | 1.06e-03 | 2.65e-03 (3.41e-04) | **8.98e-06 (7.29e-06)** |

Next, we qualitatively assess the relative power of the compared regression approaches. In the absence of known ground truth, we rely on a strong assumption that identified associations are true positives based on prior knowledge about neurobehavioral functions (Forkel et al., 2022). Therefore, we expect that many of the associations that we study in this paper should be of statistical significance. Table 3 summarizes the overall results of the illustrative study based on all three methods, providing the statistical significance of regression coefficients for all the associations that we investigate. In the table, an association where there is at least one significant regression coefficient is labeled with an asterisk. Under the strong assumption that the identified associations are true positives based on prior knowledge (Forkel et al., 2022), Table 3 can give insight into a method’s statistical power, the ability to correctly reject a null hypothesis that there is no association between a scalar factor and a white matter tract (Cohen, 2013). For most of the experiments, the results suggest that the FA Mean Regression and FMQ Regression are equally powerful methods for identifying significance, while the AFQ Regression is not a powerful method comparatively. We note that, for all AFQ and FMQ models, appropriate multiple comparisons correction using the BH-FDR method was applied to account for the multiple bundle locations/quantiles studied (as described in Sections 2.2.3 and 2.3.3). The significant AFQ and FMQ results reported in this paper have all been corrected for multiple comparisons.

**Table 3.**
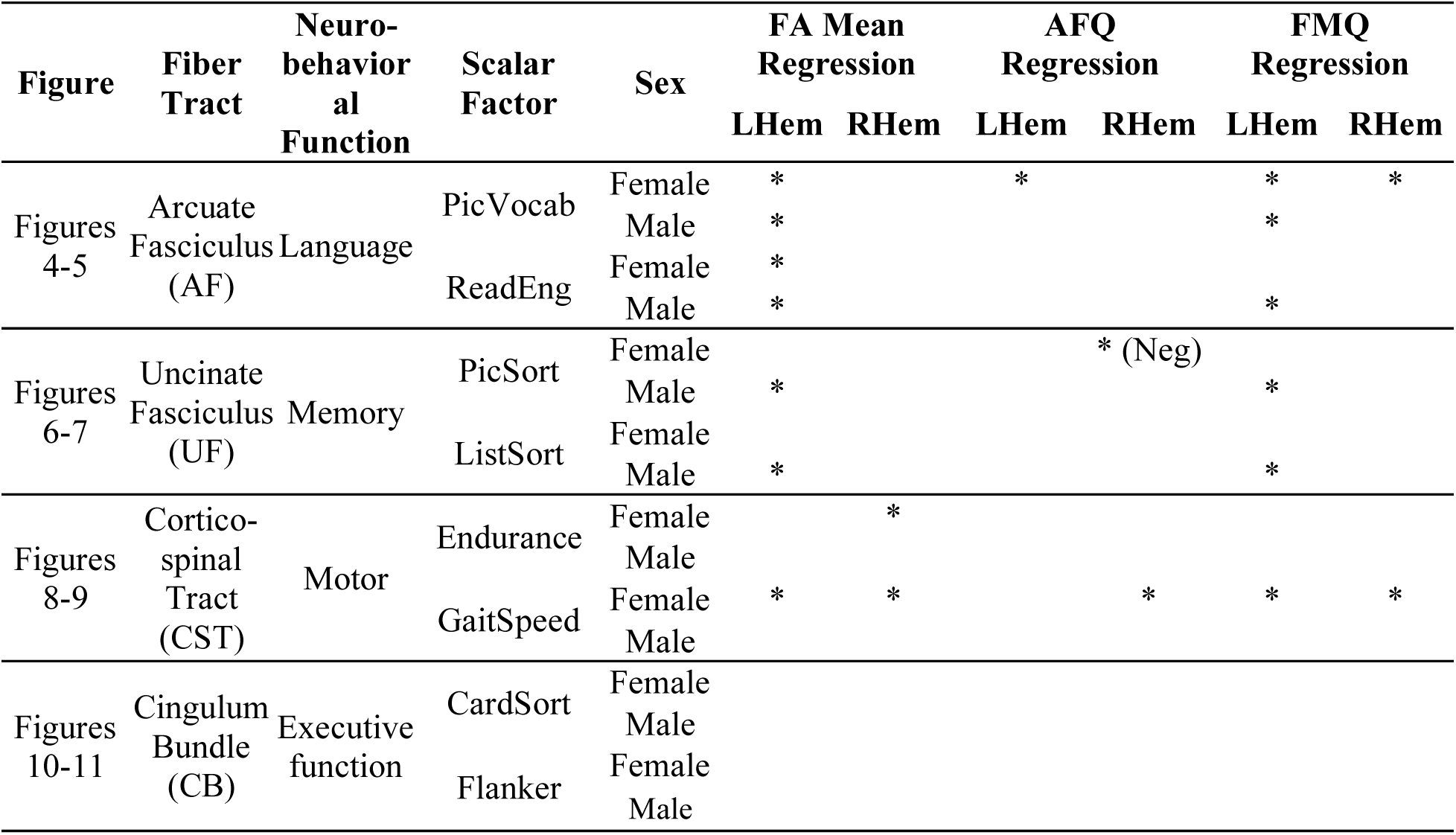
The statistical significance of regression coefficients related to the scalar factors for all the associations that we investigate. An association where there is at least one significant regression coefficient will be labeled with an asterisk. Significant negative associations are noted as *(Neg)*.

Next, we provide more detailed insight into the overall results in Table 3 by providing visualizations of the bundle experiments in Figures 4-11. The visualizations include the values of T-scores and Z-scores and how they relate to the anatomy of the studied bundles. In these figures, differences can be observed in the results of the two presumed most statistically powerful methods, FA Mean Regression and FMQ Regression. While these two methods are apparently similarly powerful in identifying associations (Table 3), the proposed FMQ Regression additionally provides the Z-scores from the periphery to the core of each bundle in both males and females (Figures 4-11). FMQ Regression therefore provides additional insight into potential anatomical underpinnings of brain-behavior associations and their differences related to sex. It can also be observed that the AFQ Regression is apparently less powerful in identifying associations.

**Figure 4:**
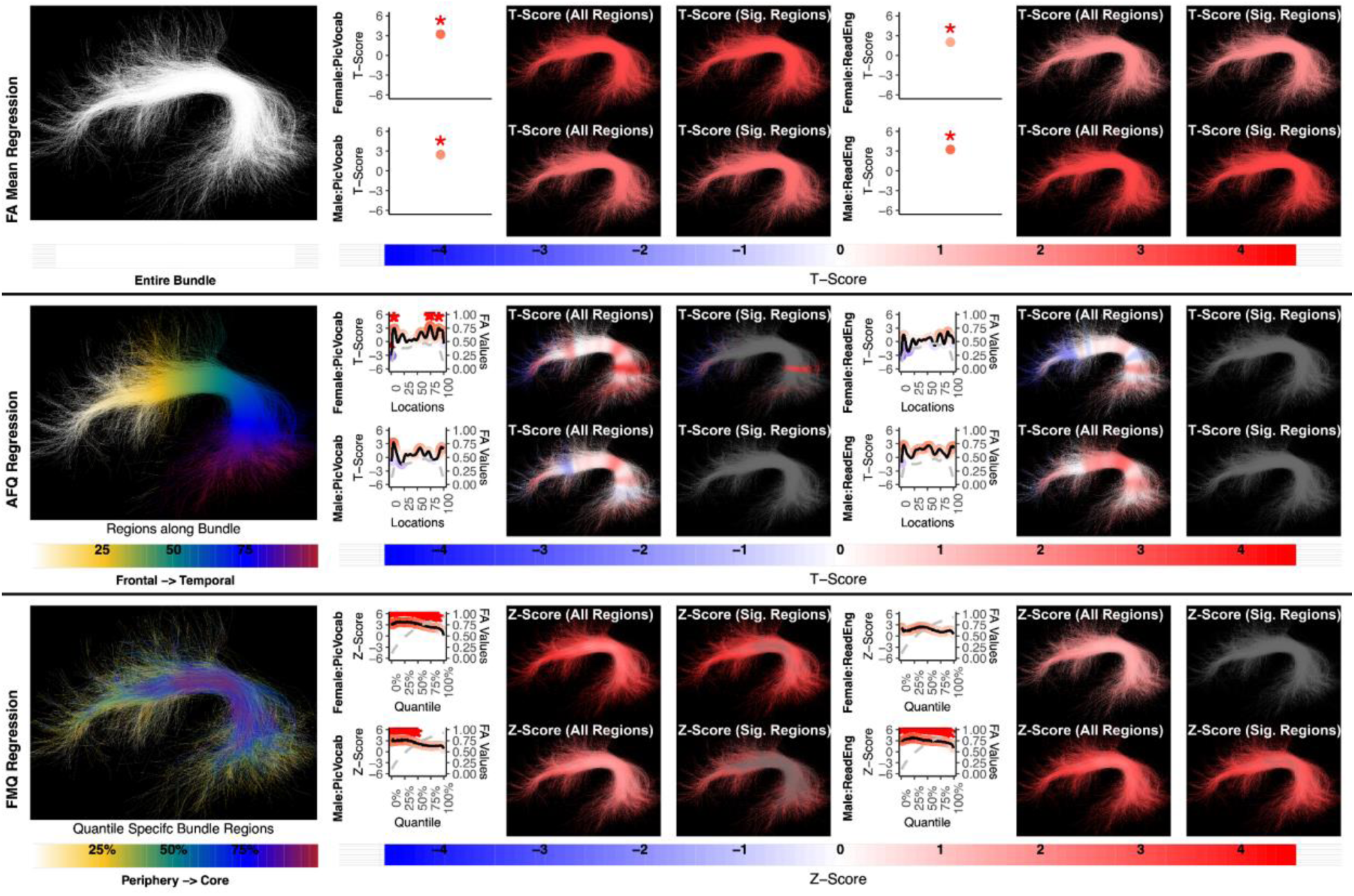
The association between AF left and language performance. FA Mean Regression identifies a significant association in PicVocab and ReadEng for males and females; AFQ Regression identifies a significant association in PicVocab in females; FMQ Regression identifies a significant association in PicVocab for males and females and ReadEng for males. Studied bundle regions are shown at left. For each experiment, plots of Z- or T-scores (solid line) and FA (dashed line) are provided, with red asterisks indicating BH-FDR-corrected statistical significance. Visualizations of Z- and T-scores are provided.

Here, we provide more details about the specific quantiles that are significant when using FMQ Regression. First, we summarize the left AF (Figure 4). Significant associations are identified for PicVocab within the left AF. In females, the quantiles from 0.5% to 90.5% are significant, covering the peripheral, intermediate, and near-core bundle regions. In males, the significant quantiles range from 0.5% to 49.5%, including peripheral and intermediate bundle regions. Additionally, FMQ Regression identifies significant associations within the left AF for ReadEng (Figure 4) for males, with significant quantiles from 0.5% to 95.5%, encompassing almost all bundle regions. Next, we summarize the right AF (Figure 5). Significant associations are identified for PicVocab within the right AF in females between quantiles 4.5% and 40.5%, covering peripheral and intermediate bundle regions. Next, we summarize the left UF (Figure 6). FMQ Regression also identifies significant associations within the left UF for PicSeq (Figure 6) for males, where quantiles 16.5% to 91.5% are significant, including near-peripheral, intermediate, and near-core bundle regions. Another significant association is observed within the left UF for ListSort (Figure 6) in males between quantiles 54.5% and 80.5%, covering the intermediate and near-core bundle regions.

**Figure 5:**
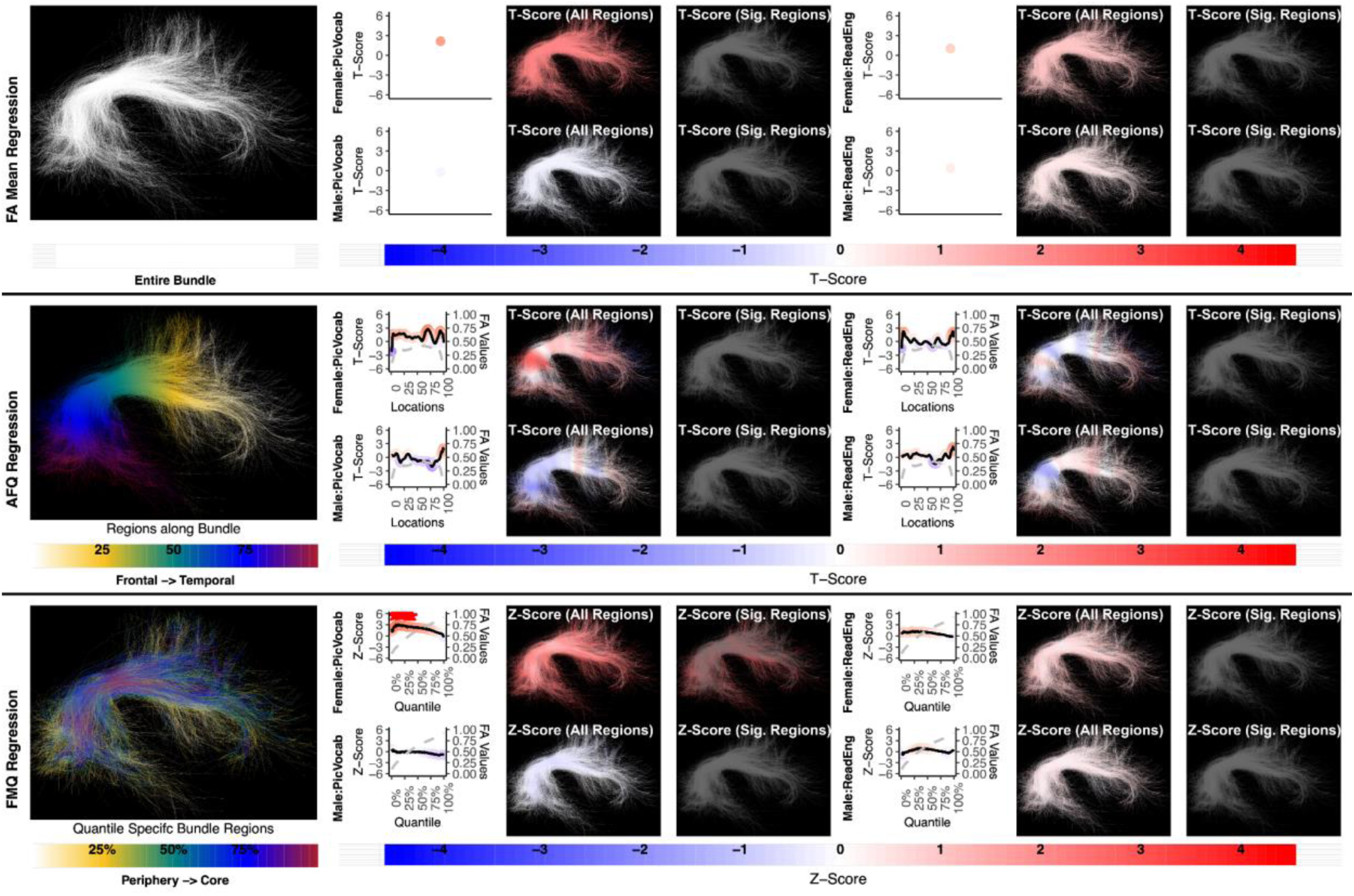
The association between AF right and language performance. FMQ Regression identifies a significant association in PicVocab for females. FA Mean Regression and AFQ Regression do not identify any statistically significant association. Studied bundle regions are shown at left. For each experiment, plots of Z- or T-scores (solid line) and FA (dashed line) are provided, with red asterisks indicating BH-FDR-corrected statistical significance. Visualizations of Z- and T-scores are provided.

**Figure 6:**
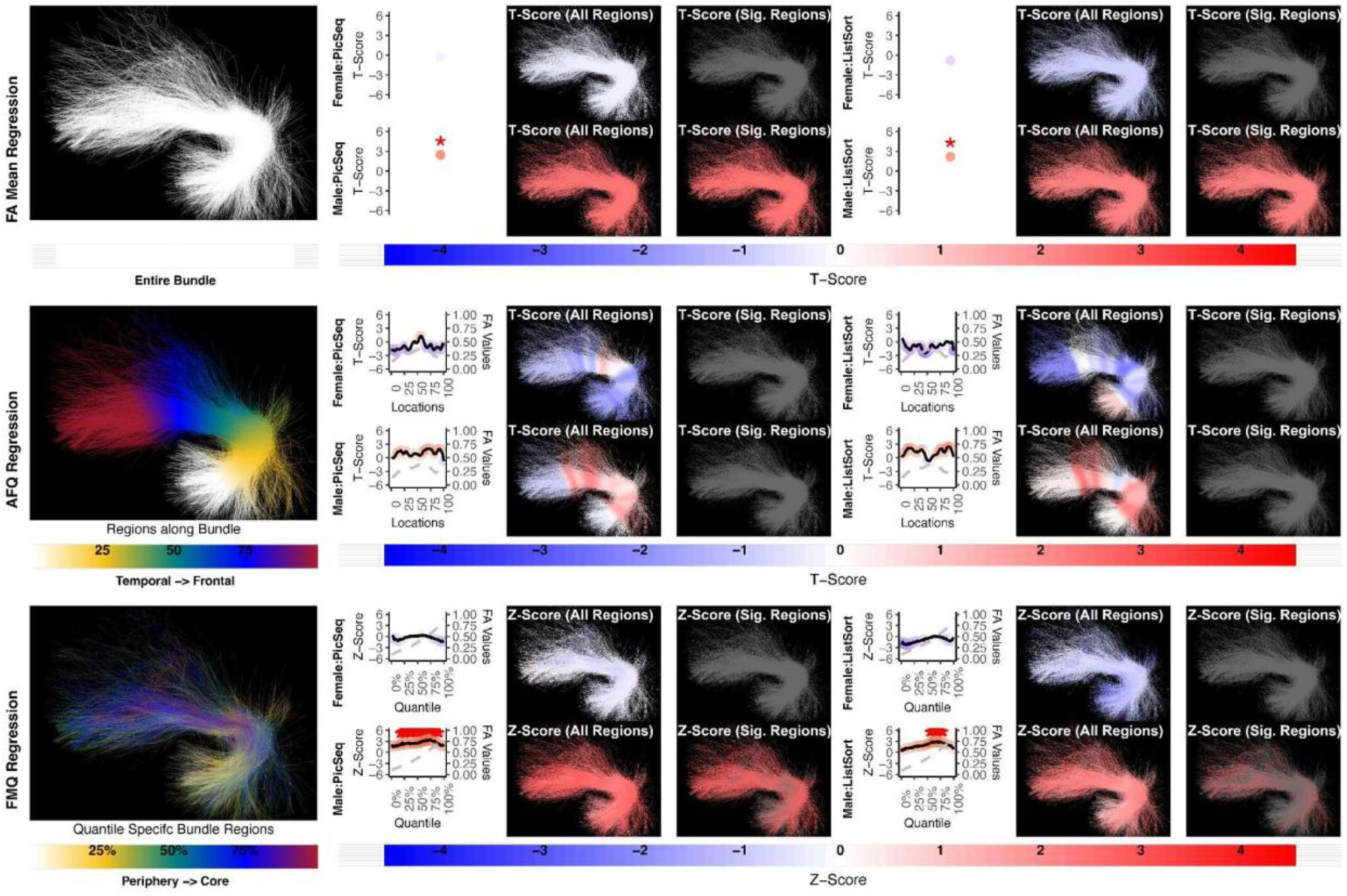
The association between UF left and memory performance. Both FA Mean Regression and FMQ Regression identify significant associations in PicSeq and ListSort for males. AFQ Regression does not identify any statistically significant association. Studied bundle regions are shown at left. For each experiment, plots of Z- or T-scores (solid line) and FA (dashed line) are provided, with red asterisks indicating BH-FDR-corrected statistical significance. Visualizations of Z- and T-scores are provided.

**Figure 7:**
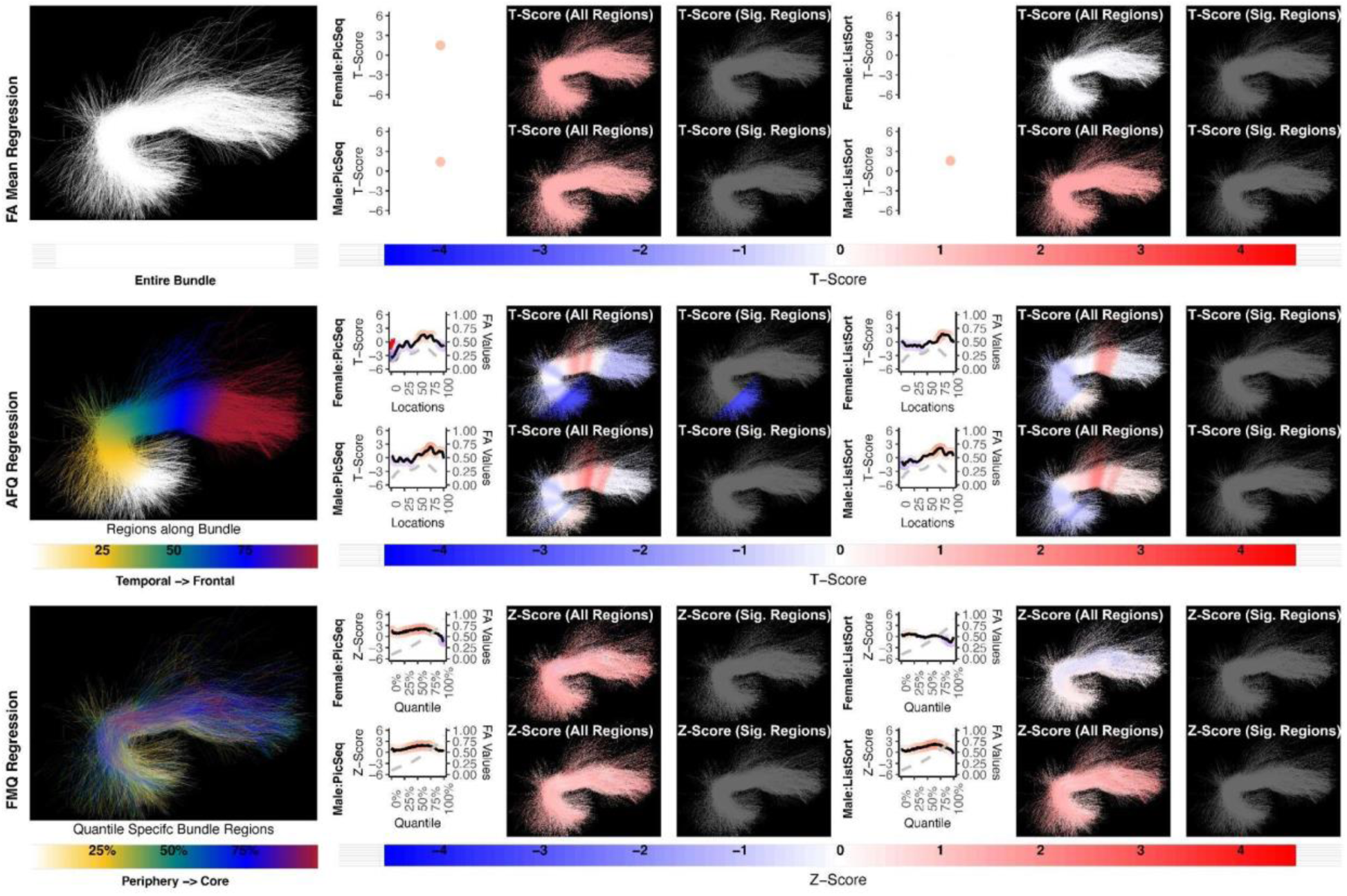
The association between UF right and memory performance. AFQ Regression identifies a negative significant association in PicSeq for males. Other methods do not identify any statistically significant association. Studied bundle regions are shown at left. Studied bundle regions are shown at left. For each experiment, plots of Z- or T-scores (solid line) and FA (dashed line) are provided, with red asterisks indicating BH-FDR-corrected statistical significance. Visualizations of Z- and T-scores are provided.

**Figure 8:**
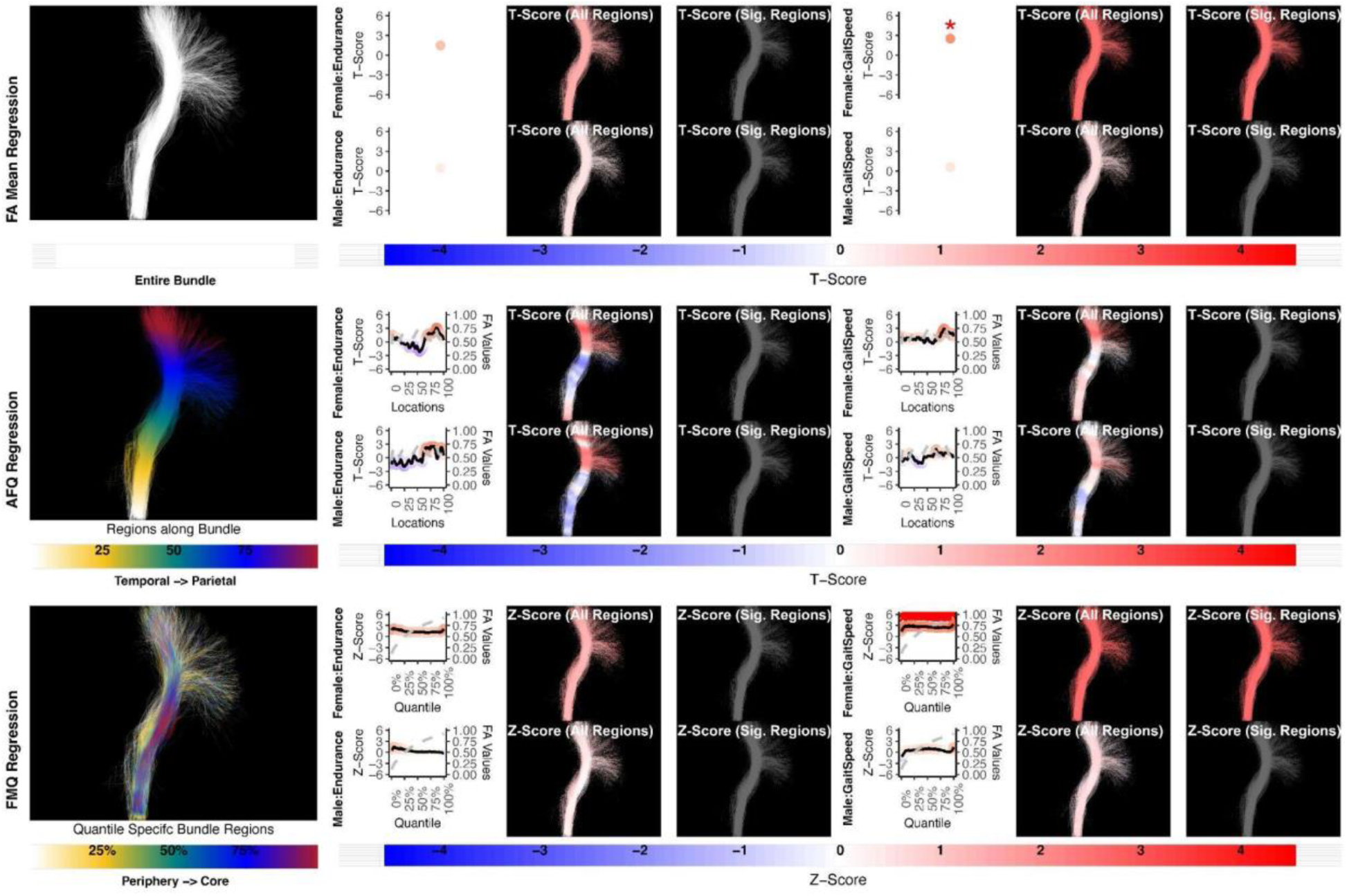
The association between CST left and motor performance. Both FA Mean Regression and FMQ Regression identify significant associations in GaitSpeed for females. AFQ Regression does not identify any statistically significant association. Studied bundle regions are shown at left. For each experiment, plots of Z- or T-scores (solid line) and FA (dashed line) are provided, with red asterisks indicating BH-FDR-corrected statistical significance. Visualizations of Z- and T-scores are provided.

**Figure 9:**
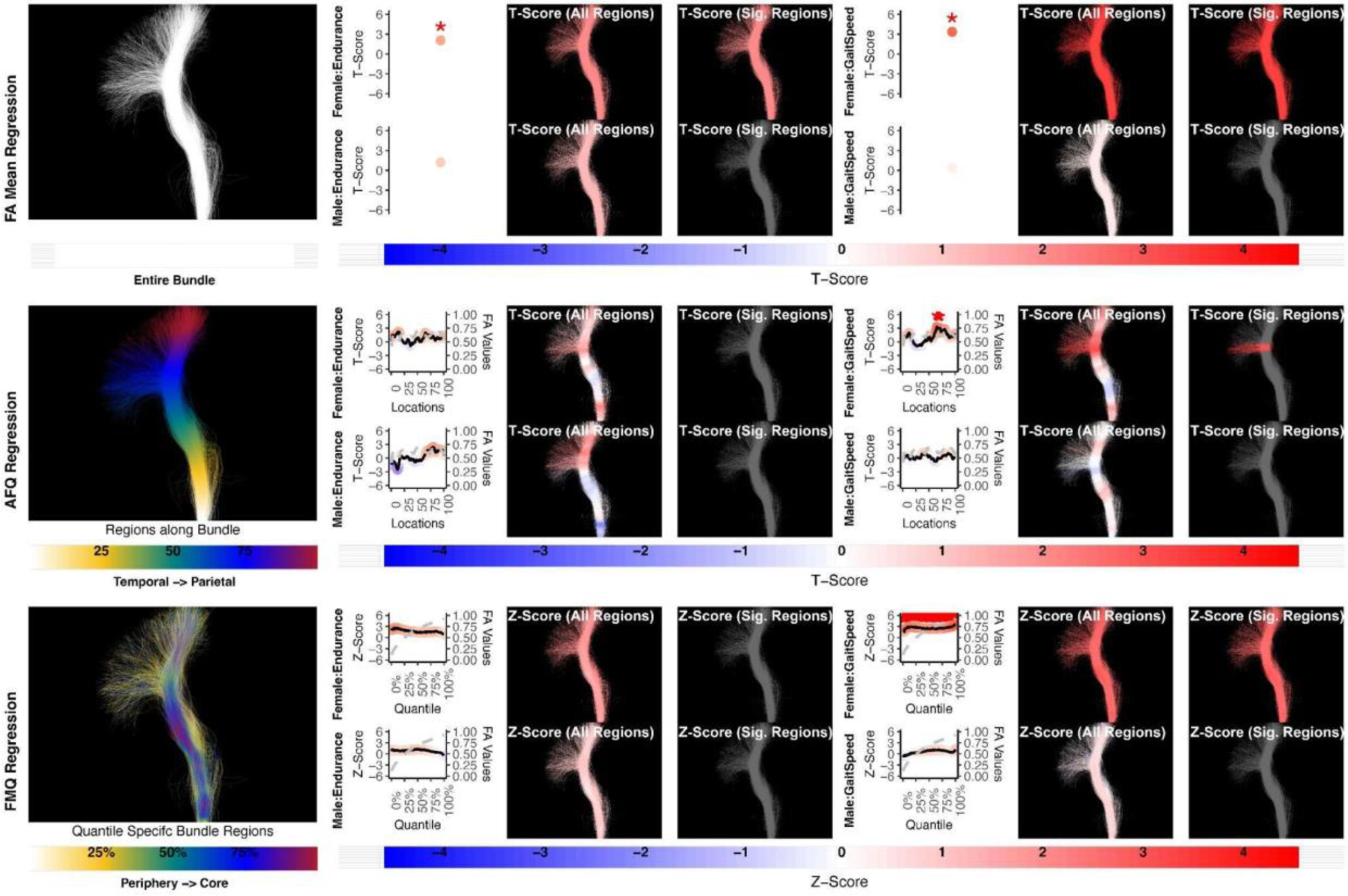
The association between CST right and motor performance. FA Mean Regression, AFQ Regression, and FMQ Regression identify significant associations in GaitSpeed for females. FA Mean Regression identifies a significant association in Endurance in females. Studied bundle regions are shown at left. Studied bundle regions are shown at left. For each experiment, plots of Z- or T-scores (solid line) and FA (dashed line) are provided, with red asterisks indicating BH-FDR-corrected statistical significance. Visualizations of Z- and T-scores are provided.

**Figure 10:**
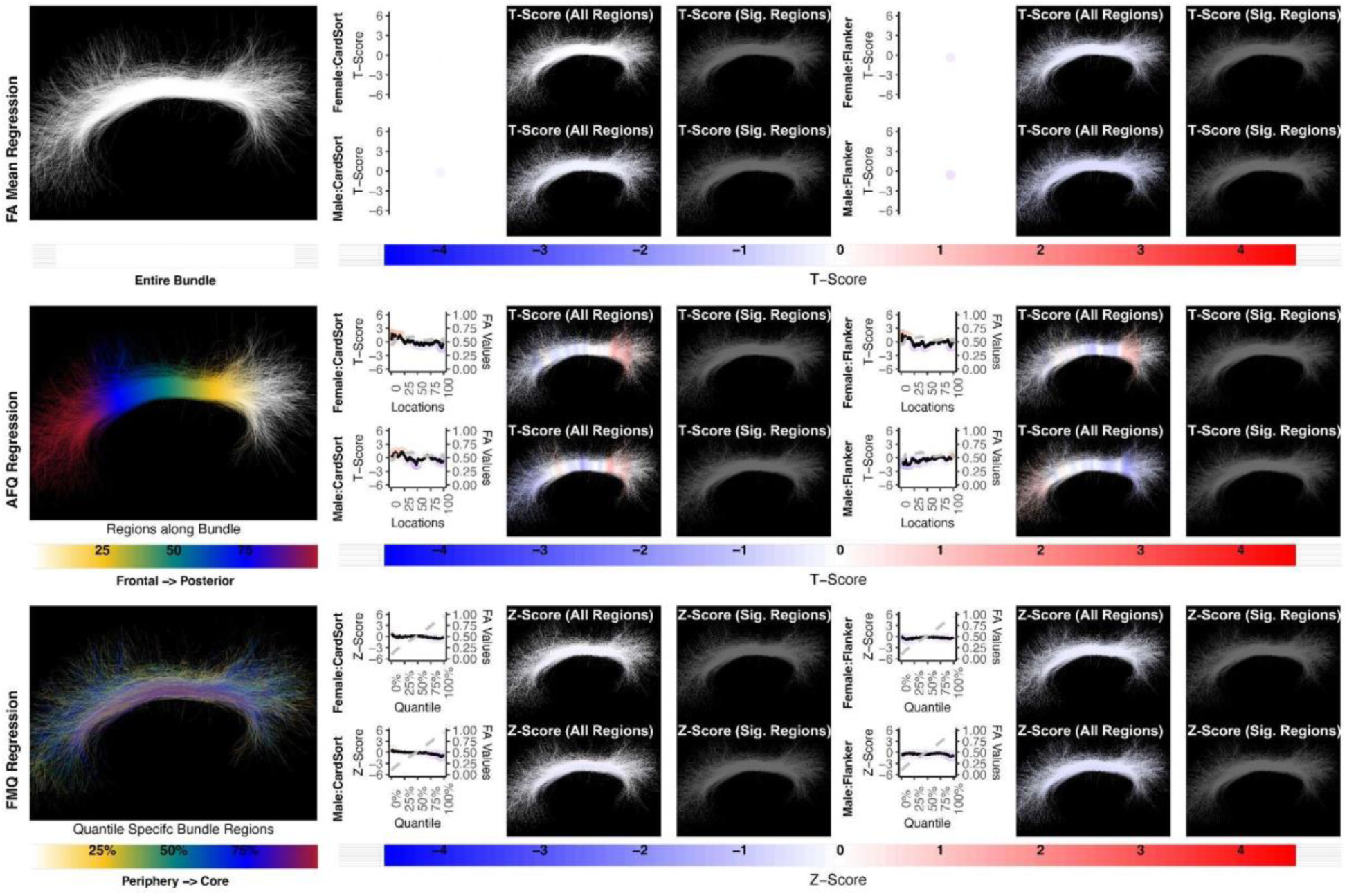
The association between CB left and executive function performance. None of the methods identify a statistically significant association. Studied bundle regions are shown at left. For each experiment, plots of Z- or T-scores (solid line) and FA (dashed line) are provided, with red asterisks indicating BH-FDR-corrected statistical significance. Studied bundle regions are shown at left. For each experiment, plots of Z- or T-scores (solid line) and FA (dashed line) are provided, with red asterisks indicating BH-FDR-corrected statistical significance. Visualizations of Z- and T-scores are provided.

**Figure 11:**
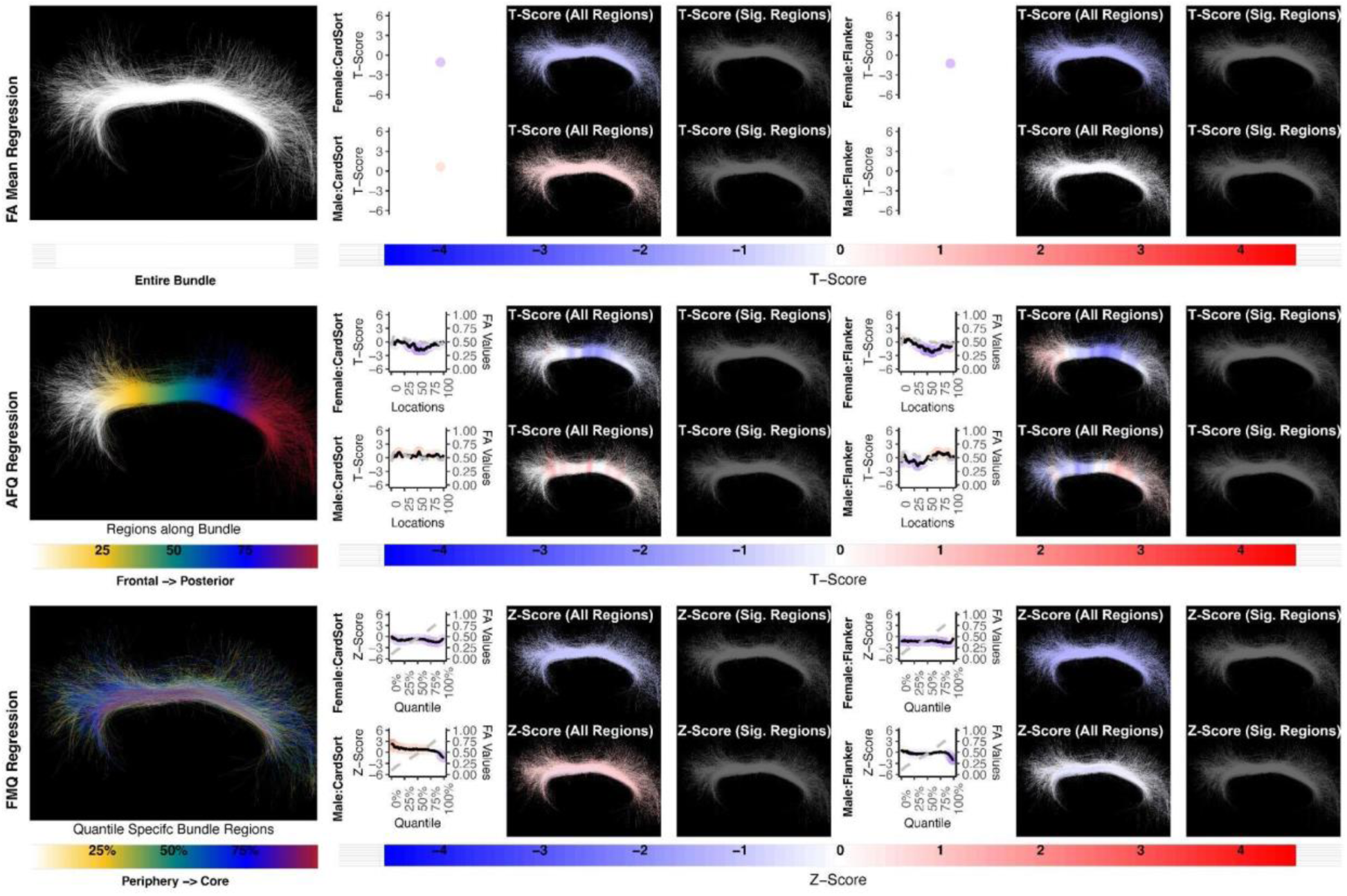
The association between CB right and executive function performance. None of the methods identify a statistically significant association. Studied bundle regions are shown at left. For each experiment, plots of Z- or T-scores (solid line) and FA (dashed line) are provided, with red asterisks indicating BH-FDR-corrected statistical significance. Studied bundle regions are shown at left. For each experiment, plots of Z- or T-scores (solid line) and FA (dashed line) are provided, with red asterisks indicating BH-FDR-corrected statistical significance. Visualizations of Z- and T-scores are provided.

Finally, we summarize the left and right CST (Figures 8 and 9). FMQ Regression identifies significant associations within the left CST for GaitSpeed (Figure 8) in females, with quantiles ranging from 2.5% to 99.5%, covering almost all bundle regions from periphery to core. Similarly, FMQ Regression identifies significant associations within the right CST for GaitSpeed in females across quantiles 0.5% to 99.5%, again covering almost all bundle regions.

## 4. Discussion

### 4.1 Methodological Contribution

In this paper, we propose a new statistical approach, FMQ Regression, for the analysis of brain fiber tract data, and we compare results to two other popular methods from the literature. We apply these methods to an illustrative study motivated by a recent review paper that describes neurobehavioral functions associated with fiber tracts in health and disease (Forkel et al., 2022). Thus, we had expected that the associations that we study in this paper would be of statistical significance. Therefore, we suggest that a method that can better identify the significance should be considered a better method. From this perspective, we make several observations. The proposed FMQ Regression generally outperforms the compared methods and produces several significant results. Our FMQ Regression results include significant findings in bundle cores (e.g., male UF bundle core FA associates with two memory performance assessments) as well as bundle peripheries (e.g., female AF periphery FA associates with PicVocab). These results motivate the potential importance of analyzing FA based on quantile-specific bundle regions, which tend to occupy regions from bundle periphery to bundle core.

We note that our method can also be applied to other microstructure measures, such as MD (which we demonstrate in Section S1 in the Supplementary Materials). Our MD results are broadly in line with our FA results, as follows. First, the supplementary investigation of MD demonstrates that FMQ Regression consistently produces the lowest MSE values compared to both MD Mean Regression and AFQ Regression models. This finding indicates that the FMQ Regression method provides a superior model fit when studying MD. Second, the FMQ Regression investigation of MD identifies multiple significant associations, including findings unique to males or females. These results suggest the general suitability of FMQ Regression for studying various measures of bundle microstructure.

In comparison with the traditional AFQ Regression strategy, a popular microstructural analysis tool, our results suggest that FMQ Regression is more powerful. For example, FMQ identifies significant associations in AF, CST, and UF, while the other two methods fail to achieve equivalent power. While FMQ Regression and AFQ Regression are both methods that provide microstructural inference, the two methods have essential differences. One difference is that they provide different approaches for region-specific analysis: bundle periphery to bundle core versus along the bundle. Another difference between the methods is that the quantile-specific bundle regions are defined in a population-based fashion, while the AFQ profile is defined in an individual-specific fashion, followed by matching across subjects. The quantile-specific bundle regions thus have the potential to reduce the effect of subject-specific sources of variability that affect the AFQ profile, such as bundle anatomical variability in shape or length, as well as FA variability in each bundle region within and across subjects. Other works have mentioned limited statistical power when using the AFQ method, which was attributed to the challenge of multiple comparisons (Richie-Halford et al., 2021). Approaches such as data reduction via feature selection (Richie-Halford et al., 2021) and suprathreshold cluster analyses (L. J. O’Donnell et al., 2009) have been proposed to reduce challenges resulting from multiple comparisons along a fiber bundle. In contrast, we note that our proposed FMQ approach has relatively high statistical power, even when using a large number of quantile-specific regions and multiple comparison adjustments. This advantage of FMQ may relate to the population-based aspect of our approach, which effectively captures variations of the data. As detecting brain-behavior associations is widely understood to be a challenging problem in neuroimaging (Gratton et al., 2022; Marek et al., 2022), new methods that can enable more powerful analysis can be a welcome addition to our toolbox.

In comparison with the traditional FA Mean Regression, the proposed FMQ Regression is similarly powerful while providing additional potential insight into associated anatomical regions (i.e., bundle core versus periphery). In Table 3, it can be seen that one significant association found by FMQ Regression was not identified by FA Mean Regression, while two significant associations identified by FA Mean Regression were not found by FMQ Regression. This suggests that the techniques are complementary, where some localized effects may be missed by the FA Mean Regression (as is well known in the literature (Colby et al., 2012; L. J. O’Donnell et al., 2009)), while some global or whole-bundle effects may be more sensitively detected by the FA Mean Regression. Overall, the fact that the significant findings were generally consistent across the FMQ and Mean FA Regression methods, across multiple NIH toolbox measures (e.g., of language function), and across hemispheres (e.g., bilateral female CST effects), suggests the robustness of the proposed FMQ Regression.

Our FMQ Regression is different from other fiber tract data analyses using the quantile regression technique (Lv et al., 2021; Ryan et al., 2022). In contrast to our approach, the previous works (Lv et al., 2021; Ryan et al., 2022) rely on the FA mean of the entire fiber tract (measured at the individual level) and investigate the association between the conditional quantile of the FA mean and covariates of interest. We also note that the proposed FMQ Regression is a population-based analysis and inherently differs from an individual-level analysis (which could be feasible by calculating the quantiles of each individual’s fiber tract). Such an individual-level analysis would not rely on the methodology of quantile regression but on an ordinary regression method (e.g., ordinary least squares regression). We further note that ordinary regression assumes the error distribution is normal. In contrast, quantile regression does not make such strict assumptions and is, therefore, more robust (Wei et al., 2006).

### 4.2 Neuroscience Findings

In the following paragraphs, we briefly discuss our current findings in each fiber tract in relation to the literature.

Consistent with many reports of associations between the left, but not right, AF and language performance in healthy individuals (Yeatman et al., 2011; Zekelman et al., 2022), the proposed FMQ Regression identified a statistically significant association in the left AF in both males and females. However, in females only, the FMQ Regression (and not the FA mean or AFQ methods) identified a statistically significant association with PicVocab in the bundle periphery and intermediate bundle regions of the right AF. This is of interest for further investigation and could potentially relate to known sex differences in AF, such as its greater symmetry in females (Thiebaut de Schotten et al., 2011). Interestingly, our significant findings relating the left AF FA to language performance spanned many quantiles of FA but never included the maximum FA bundle “core” regions (i.e., the highest quantiles near 100% were never significant, as shown in Figure 4). In fact, the relationships between AF microstructure (FA) and two assessments of language are stronger in the periphery and decrease toward the core of the bundle in both males and females. This observation may potentially represent a challenge for uncovering brain-behavior language associations using TBSS (tract-based spatial statistics), a method that focuses only on maximum-FA voxels thought to represent bundle cores (Smith et al., 2006). For example, a recent investigation studied six different language assessments using TBSS and found only one association of FA in the left superior longitudinal fasciculus, which includes the AF (Houston et al., 2019). Converging evidence from our recent geometric machine learning work also identified peripheral regions of AF, including regions of the gray-white matter interface, to be most predictive of individual performance on language assessments (Chen et al., 2024). This is in line with recent work investigating the shape of the white matter association tracts (including AF), which shows that the peripheral regions where bundles originate and terminate in the cortex have a large degree of inter-individual variability and are therefore a good descriptor of inter-individual differences in white matter structure (Yeh, 2020).

Consistent with a handful of other studies of UF in healthy individuals (Mabbott et al., 2009; Schaeffer et al., 2014), the proposed FMQ Regression identified a statistically significant association between left UF FA and memory performance (Figure 6). However, in the present study, this effect was observed only in males (in intermediate and near-core bundle regions). This finding motivates the importance of studying sex effects in the relationship between brain microstructure and individual functional performance.

While it is well understood that the CST subserves motor function (Welniarz et al., 2017) and is left-lateralized (Thiebaut de Schotten et al., 2011), most existing studies of FA and motor function have been performed in patients with diminished function. However, a recent study in healthy young adults showed that CST FA was bilaterally associated with corticospinal excitability, a transcranial magnetic stimulation measure of individual function (Betti et al., 2022). Our finding that CST FA is associated with motor functional performance (GaitSpeed) bilaterally, and only in females, further motivates the need to study tract microstructure and its relationship to human brain functional performance in both males and females, as well as in healthy individuals (Fig. 7-8).

We report negative results (no significance) for the relationship between CB FA and measures of executive function (CardSort and Flanker) in healthy young adults using all compared regression methods (Fig. 9-10). A recent review on white matter tracts and executive function suggests a role for CB, especially in inhibition; however, the supporting neuroimaging studies include aging and neuropsychiatric populations, not healthy individuals (Ribeiro et al., 2024). Our results do not contradict the potential role of CB in executive function in healthy young adults; our findings merely indicate that CB FA microstructure does not relate to executive function performance in the study population. In this study, we focused on the superior part of the CB, excluding the temporal portion of the bundle, because individual streamlines generally do not trace the entire trajectory of the CB. This is in part because axons enter and leave the CB along its entire length (Heilbronner & Haber, 2014; Jones et al., 2013). In addition, the superior part of the cingulum more closely correlates with attention and executive function, whereas the temporal cingulum is associated with episodic memory (Kantarci et al., 2011; Metzler-Baddeley et al., 2012).

### 4.3 Neuroanatomical Discussion

In this paper, we have proposed analyzing white matter bundles in dMRI using regions that are defined using quantiles of FA, with the result that the quantile-specific regions are approximately defined from the bundle periphery to the bundle core. Neuroanatomical research demonstrates that for many bundles, axons enter and leave the bundle along its course (Heilbronner & Haber, 2014; Morris et al., 1999; Mufson & Pandya, 1984; Petrides & Pandya, 2007; Schmahmann & Pandya, 2009; Yakovlev & Locke, 1961). As these axons leave the bundle, they curve and necessarily intersect and cross axons from other bundles closer to their cortical or subcortical targets. This intersection will lead to a lower FA (Jones, 2010). In the current study, these regions of lower FA are located toward the ends of the tract as the streamlines fan out as a spray of fibers and traverse the periphery of other fiber bundles (Makris et al., 1997). In addition, these regions of lower FA in peripheral locations are also observed along the length of the fiber, which corresponds to anatomical studies of the cingulum bundle that show that axons enter or leave the tract along its course (Mufson & Pandya, 1984; Yakovlev & Locke, 1961).

### 4.3 Limitations and Future Work

Potential limitations of the present study, including suggested future work to address limitations, are as follows.

In this work, we demonstrated the estimation of *C* = 100 sets of coefficients for all the quantile-specific bundle regions. Future work may investigate other numbers of quantile-specific bundle regions by varying *C*. From the computational perspective, our method requires a large streamline dataset sampled from all subject fiber bundles under study in the population. Future work may investigate optimizing the amount of input data needed to obtain results in very large datasets.

Our current paper focused on FA, with supplementary results showing the application of FMQ to study MD. In the future, it will be of interest to investigate the application of FMQ Regression, potentially in a multivariate fashion, to study multiple measures of microstructure or imaging data within fiber tracts.

In this study, we have described quantile locations as bundle core, intermediate bundle regions, or bundle periphery regions, with accompanying visualizations to provide more details. However, it can be observed that not all fiber bundles are completely included in the field of view (FOV) of a dMRI scan, which can make the periphery-core interpretation more nuanced (Chen et al., 2025). For instance, the high-FA “core” region of the CST can be observed to continue inferiorly in the brainstem, which extends outside of the FOV. Another consideration is the potential presence of somatotopic or functional subdivisions of a fiber bundle. For instance, future work could separately study associations of individual bundles within the CST, e.g., those originating in trunk, leg, hand, and face motor cortical regions (He et al., 2023) or bundles representing subdivisions within the AF (Fernández-Miranda et al., 2015). The current study is a fiber-bundle-based study. Another possible future work might involve a complete whole-brain tractography as the input, which could enable the investigation of microstructure quantiles in the entire white matter.

In this initial paper describing the proposed FMQ Regression method, we have performed a testbed study of selected white matter fiber bundles, where we focused on one selected microstructure measure (FA) and studied multiple selected neurobehavioral measures. In future work, it will be interesting to extend and apply the proposed FMQ approach to perform studies of additional datasets, fiber bundles, microstructure measures, and scalar factors of interest, potentially leading to deeper insights into the brain’s structural-functional relationships.

## 5. Conclusion

We have proposed FMQ Regression, a novel quantile regression methodology for studying white matter bundles in the brain. We find that analyzing FA using quantile-specific bundle regions, which tend to define regions from bundle periphery to bundle core, is much more powerful than a traditional AFQ method that spatially subdivides bundles along their lengths. Our results suggest that FMQ Regression is a powerful tool for studying brain-behavior associations using white matter tractography data.

## Data Availability

This study utilized publicly available data from HCP-YA (https://www.humanconnectome.org/study/hcp-young-adult/overview). Data can be accessed via data use agreements.

## Author Contributions

**Zhou Lan**: Visualization, Conceptualization, Formal analysis, Writing – original draft, Writing – review & editing. **Yuqian Chen**: Formal analysis, Writing – review & editing. **Jarrett Rushmore**: Formal analysis, Writing – review & editing. **Leo Zekelman:** Writing – review & editing. **Nikos Makris:** Writing – review & editing. **Yogesh Rathi:** Writing – review & editing. **Alexandra J. Golby:** Writing – review & editing. **Fan Zhang:** Formal analysis, Writing – review & editing. **Lauren J. O’Donnell**: Visualization, Conceptualization, Writing – original draft, Writing – review & editing.

## Acknowledgments

We gratefully acknowledge funding provided by the following grants: National Institutes of Health (NIH) grants R01MH132610, R01MH125860, R01MH119222, R01NS125307, R01NS125781, R21NS136960, R01AG042512, R01DC020965.

## Footnotes

1 https://workshop.dipy.org/documentation/0.16.0./examples_built/afq_tract_profiles/

## References

Basser, P. J. (1995). Inferring microstructural features and the physiological state of tissues from diffusion-weighted images. NMR in Biomedicine, 8(7-8), 333–344.

Basser, P. J., Pajevic, S., Pierpaoli, C., Duda, J., & Aldroubi, A. (2000). In vivo fiber tractography using DT-MRI data. Magnetic Resonance in Medicine: Official Journal of the Society of Magnetic Resonance in Medicine / Society of Magnetic Resonance in Medicine, 44(4), 625–632.

Basser, P. J., & Pierpaoli, C. (2011). Microstructural and physiological features of tissues elucidated by quantitative-diffusion-tensor MRI. Journal of Magnetic Resonance, 213(2), 560–570.

Batchelor, P. G., Calamante, F., Tournier, J.-D., Atkinson, D., Hill, D. L. G., & Connelly, A. (2006). Quantification of the shape of fiber tracts. Magnetic Resonance in Medicine, 55(4), 894–903.

Benjamini, Y., & Yekutieli, D. (2001). The control of the false discovery rate in multiple testing under dependency. Annals of Statistics, 29(4), 1165–1188.

Betti, S., Fedele, M., Castiello, U., Sartori, L., & Budisavljević, S. (2022). Corticospinal excitability and conductivity are related to the anatomy of the corticospinal tract. Brain Structure & Function, 227(3), 1155–1164.

Bozzali, M., Falini, A., Franceschi, M., Cercignani, M., Zuffi, M., Scotti, G., Comi, G., & Filippi, M. (2002). White matter damage in Alzheimer’s disease assessed in vivo using diffusion tensor magnetic resonance imaging. *Journal of Neurology*, Neurosurgery, and Psychiatry, 72(6), 742–746.

Catani, M., Howard, R. J., Pajevic, S., & Jones, D. K. (2002). Virtual in vivo interactive dissection of white matter fasciculi in the human brain. NeuroImage, 17(1), 77–94.

Cetin-Karayumak, S., Zhang, F., Zurrin, R., Billah, T., Zekelman, L., Makris, N., Pieper, S., O’Donnell, L. J., & Rathi, Y. (2024). Harmonized diffusion MRI data and white matter measures from the Adolescent Brain Cognitive Development Study. Scientific Data, 11(1), 249.

Chamberland, M., St-Jean, S., Tax, C. M. W., & Jones, D. K. (2019). Obtaining representative core streamlines for white matter tractometry of the human brain. In Computational Diffusion MRI (pp. 359–366). Springer International Publishing.

Chandio, B. Q., Risacher, S. L., Pestilli, F., Bullock, D., Yeh, F.-C., Koudoro, S., Rokem, A., Harezlak, J., & Garyfallidis, E. (2020). Bundle analytics, a computational framework for investigating the shapes and profiles of brain pathways across populations. Scientific Reports, 10(1), 17149.

Chenot, Q., Tzourio-Mazoyer, N., Rheault, F., Descoteaux, M., Crivello, F., Zago, L., Mellet, E., Jobard, G., Joliot, M., Mazoyer, B., & Petit, L. (2019). A population-based atlas of the human pyramidal tract in 410 healthy participants. Brain Structure & Function, 224(2), 599–612.

Chen, Y., Zekelman, L., Lo, Y., Cetin-Karayumak, S., Xue, T., Rathi, Y., Makris, N., Zhang, F., Cai, W., & J O’donnell, L. (2025). TractCloud-FOV: Deep learning-based robust tractography parcellation in diffusion MRI with incomplete field of view. In arXiv [cs.CV]. arXiv. http://arxiv.org/abs/2502.20637

Chen, Y., Zekelman, L. R., Zhang, C., Xue, T., Song, Y., Makris, N., Rathi, Y., Golby, A. J., Cai, W., Zhang, F., & O’Donnell, L. J. (2024). TractGeoNet: A geometric deep learning framework for pointwise analysis of tract microstructure to predict language assessment performance. Medical Image Analysis, 94(103120), 103120.

Ciccarelli, O., Parker, G. J. M., Toosy, A. T., Wheeler-Kingshott, C. A. M., Barker, G. J., Boulby, P. A., Miller, D. H., & Thompson, A. J. (2003). From diffusion tractography to quantitative white matter tract measures: a reproducibility study. NeuroImage, 18(2), 348– 359.

Cohen, J. (2013). Statistical power analysis for the behavioral sciences (2nd ed.). Routledge. 10.4324/9780203771587

Colby, J. B., Soderberg, L., Lebel, C., Dinov, I. D., Thompson, P. M., & Sowell, E. R. (2012). Along-tract statistics allow for enhanced tractography analysis. NeuroImage, 59(4), 3227– 3242.

Corouge, I., Fletcher, P. T., Joshi, S., Gouttard, S., & Gerig, G. (2006). Fiber tract-oriented statistics for quantitative diffusion tensor MRI analysis. Medical Image Analysis, 10(5), 786–798.

Corouge, I., Gouttard, S., & Gerig, G. (2004). A statistical shape model of individual fiber tracts extracted from diffusion tensor MRI. In Medical Image Computing and Computer-Assisted Intervention – MICCAI 2004 (pp. 671–679). Springer Berlin Heidelberg.

Daducci, A., Canales-Rodríguez, E. J., Zhang, H., Dyrby, T. B., Alexander, D. C., & Thiran, J.-P. (2015). Accelerated Microstructure Imaging via Convex Optimization (AMICO) from diffusion MRI data. NeuroImage, 105, 32–44.

Damatac, C. G., Chauvin, R. J. M., Zwiers, M. P., van Rooij, D., Akkermans, S. E. A., Naaijen, J., Hoekstra, P. J., Hartman, C. A., Oosterlaan, J., Franke, B., Buitelaar, J. K., Beckmann, C. F., & Sprooten, E. (2022). White matter microstructure in attention-deficit/hyperactivity disorder: A systematic tractography study in 654 individuals. Biological Psychiatry: Cognitive Neuroscience and Neuroimaging, 7(10), 979–988.

Elias, G. J. B., Germann, J., Joel, S. E., Li, N., Horn, A., Boutet, A., & Lozano, A. M. (2024). A large normative connectome for exploring the tractographic correlates of focal brain interventions. Scientific Data, 11(1), 353.

Farquharson, S., Tournier, J.-D., Calamante, F., Fabinyi, G., Schneider-Kolsky, M., Jackson, G. D., & Connelly, A. (2013). White matter fiber tractography: why we need to move beyond DTI. Journal of Neurosurgery, 118(6), 1367–1377.

Fernández-Miranda, J. C., Wang, Y., Pathak, S., Stefaneau, L., Verstynen, T., & Yeh, F.-C. (2015). Asymmetry, connectivity, and segmentation of the arcuate fascicle in the human brain. Brain Structure & Function, 220(3), 1665–1680.

Fields, R. D. (2008). White matter in learning, cognition and psychiatric disorders. Trends in Neurosciences, 31(7), 361–370.

Forkel, S. J., Friedrich, P., Thiebaut de Schotten, M., & Howells, H. (2022). White matter variability, cognition, and disorders: a systematic review. Brain Structure & Function, 227(2), 529–544.

Gari, I. B., Javid, S., Zhu, A. H., Gadewar, S. P., Narula, S., Ramesh, A., Thomopoulos, S. I., Strike, L., Zubicaray, G. I. de, McMahon, K. L., Wright, M. J., Thompson, P. M., Nir, T. M., & Jahanshad, N. (2023). Along-tract parameterization of white matter microstructure using medial tractography analysis (MeTA). 2023 19th International Symposium on Medical Information Processing and Analysis (SIPAIM), 1–5.

Garyfallidis, E., Brett, M., Amirbekian, B., Rokem, A., van der Walt, S., Descoteaux, M., Nimmo-Smith, I., & Dipy Contributors. (2014). Dipy, a library for the analysis of diffusion MRI data. Frontiers in Neuroinformatics, 8, 8.

Garyfallidis, E., Brett, M., Correia, M. M., Williams, G. B., & Nimmo-Smith, I. (2012). QuickBundles, a Method for Tractography Simplification. Frontiers in Neuroscience, 6, 175.

Garyfallidis, E., Côté, M.-A., Rheault, F., Sidhu, J., Hau, J., Petit, L., Fortin, D., Cunanne, S., & Descoteaux, M. (2018). Recognition of white matter bundles using local and global streamline-based registration and clustering. NeuroImage, 170, 283–295.

Gerig, G., Gouttard, S., & Corouge, I. (2004). Analysis of brain white matter via fiber tract modeling. Conference Proceedings: … Annual International Conference of the IEEE Engineering in Medicine and Biology Society. IEEE Engineering in Medicine and Biology Society. Conference, 2004, 4421–4424.

Gershon, R. C., Cook, K. F., Mungas, D., Manly, J. J., Slotkin, J., Beaumont, J. L., & Weintraub, S. (2014). NIH Toolbox Oral Reading Recognition Test. Journal of the International Neuropsychological Society: JINS. 10.1037/t63740-000

Glasser, M. F., Sotiropoulos, S. N., Wilson, J. A., Coalson, T. S., Fischl, B., Andersson, J. L., Xu, J., Jbabdi, S., Webster, M., Polimeni, J. R., Van Essen, D. C., Jenkinson, M., & WU-Minn HCP Consortium. (2013). The minimal preprocessing pipelines for the Human Connectome Project. NeuroImage, 80, 105–124.

Gratton, C., Nelson, S. M., & Gordon, E. M. (2022). Brain-behavior correlations: Two paths toward reliability. Neuron, 110(9), 1446–1449.

Hastie, T., Tibshirani, R., & Friedman, J. (2001). The elements of statistical learning. Springer.

Heilbronner, S. R., & Haber, S. N. (2014). Frontal cortical and subcortical projections provide a basis for segmenting the cingulum bundle: implications for neuroimaging and psychiatric disorders. The Journal of Neuroscience: The Official Journal of the Society for Neuroscience, 34(30), 10041–10054.

He, J., Zhang, F., Pan, Y., Feng, Y., Rushmore, J., Torio, E., Rathi, Y., Makris, N., Kikinis, R., Golby, A. J., & O’Donnell, L. J. (2023). Reconstructing the somatotopic organization of the corticospinal tract remains a challenge for modern tractography methods. Human Brain Mapping, 44(17), 6055–6073.

Hodes, R. J., Insel, T. R., Landis, S. C., & NIH Blueprint for Neuroscience Research. (2013). The NIH toolbox: setting a standard for biomedical research. Neurology, 80(11 Suppl 3), S1.

Houston, J., Allendorfer, J., Nenert, R., Goodman, A. M., & Szaflarski, J. P. (2019). White Matter Language Pathways and Language Performance in Healthy Adults Across Ages. Frontiers in Neuroscience, 13, 1185.

Jeurissen, B., Leemans, A., Tournier, J.-D., Jones, D. K., & Sijbers, J. (2013). Investigating the prevalence of complex fiber configurations in white matter tissue with diffusion magnetic resonance imaging. Human Brain Mapping, 34(11), 2747–2766.

Johnson, C. A., Liu, Y., Waller, N., & Chang, S.-E. (2022). Tract profiles of the cerebellar peduncles in children who stutter. Brain Structure & Function, 227(5), 1773–1787.

Jones, D. K. (2010). Challenges and limitations of quantifying brain connectivityin vivowith diffusion MRI. Imaging in Medicine, 2(3), 341–355.

Jones, D. K., Christiansen, K. F., Chapman, R. J., & Aggleton, J. P. (2013). Distinct subdivisions of the cingulum bundle revealed by diffusion MRI fibre tracking: implications for neuropsychological investigations. Neuropsychologia, 51(1), 67–78.

Kantarci, K., Senjem, M. L., Avula, R., Zhang, B., Samikoglu, A. R., Weigand, S. D., Przybelski, S. A., Edmonson, H. A., Vemuri, P., Knopman, D. S., Boeve, B. F., Ivnik, R. J., Smith, G. E., Petersen, R. C., & Jack, C. R., Jr. (2011). Diffusion tensor imaging and cognitive function in older adults with no dementia. Neurology, 77(1), 26–34.

Koenker, R., Chernozhukov, V., He, X., & Peng, L. (Eds.). (2020). Handbook of quantile regression. Chapman & Hall/CRC.

Kruper, J., Benson, N. C., Caffarra, S., Owen, J., Wu, Y., Lee, A. Y., Lee, C. S., Yeatman, J. D., Rokem, A., & UK Biobank Eye and Vision Consortium. (2023). Optic radiations representing different eccentricities age differently. Human Brain Mapping, 44(8), 3123– 3135.

Kruper, J., Hagen, M. P., Rheault, F., Crane, I., Gilmore, A., Narayan, M., Motwani, K., Lila, E., Rorden, C., Yeatman, J. D., & Rokem, A. (2024). Tractometry of the Human Connectome Project: resources and insights. Frontiers in Neuroscience, 18, 1389680.

Loring, D. W., Bowden, S. C., Staikova, E., Bishop, J. A., Drane, D. L., & Goldstein, F. C. (2019). NIH Toolbox Picture Sequence Memory Test for Assessing Clinical Memory Function: Diagnostic Relationship to the Rey Auditory Verbal Learning Test. Archives of Clinical Neuropsychology: The Official Journal of the National Academy of Neuropsychologists, 34(2), 268–276.

Lv, J., Di Biase, M., Cash, R. F. H., Cocchi, L., Cropley, V. L., Klauser, P., Tian, Y., Bayer, J., Schmaal, L., Cetin-Karayumak, S., Rathi, Y., Pasternak, O., Bousman, C., Pantelis, C., Calamante, F., & Zalesky, A. (2021). Individual deviations from normative models of brain structure in a large cross-sectional schizophrenia cohort. Molecular Psychiatry, 26(7), 3512– 3523.

Mabbott, D. J., Rovet, J., Noseworthy, M. D., Smith, M. L., & Rockel, C. (2009). The relations between white matter and declarative memory in older children and adolescents. Brain Research, 1294, 80–90.

Maddah, M., Grimson, W. E. L., & Warfield, S. K. (2006). Statistical modeling and EM clustering of white matter fiber tracts. 3rd IEEE International Symposium on Biomedical Imaging: Macro to Nano, 2006. 3rd IEEE International Symposium on Biomedical Imaging: Macro to Nano, 2006., Arlington, Virginia, USA. 10.1109/isbi.2006.1624850

Makris, N., Worth, A. J., Papadimitriou, G. M., Stakes, J. W., Caviness, V. S., Kennedy, D. N., Pandya, D. N., Kaplan, E., Sorensen, A. G., Wu, O., Reese, T. G., Wedeen, V. J., Rosen, B. R., Kennedy, D. N., & Davis, T. L. (1997). Morphometry of in vivo human white matter association pathways with diffusion-weighted magnetic resonance imaging. Annals of Neurology, 42(6), 951–962.

Marek, S., Tervo-Clemmens, B., Calabro, F. J., Montez, D. F., Kay, B. P., Hatoum, A. S., Donohue, M. R., Foran, W., Miller, R. L., Hendrickson, T. J., Malone, S. M., Kandala, S., Feczko, E., Miranda-Dominguez, O., Graham, A. M., Earl, E. A., Perrone, A. J., Cordova, M., Doyle, O., … Dosenbach, N. U. F. (2022). Reproducible brain-wide association studies require thousands of individuals. Nature, 603(7902), 654–660.

Metzler-Baddeley, C., Jones, D. K., Steventon, J., Westacott, L., Aggleton, J. P., & O’Sullivan, M. J. (2012). Cingulum microstructure predicts cognitive control in older age and mild cognitive impairment. The Journal of Neuroscience: The Official Journal of the Society for Neuroscience, 32(49), 17612–17619.

Morris, R., Pandya, D. N., & Petrides, M. (1999). Fiber system linking the mid-dorsolateral frontal cortex with the retrosplenial/presubicular region in the rhesus monkey. The Journal of Comparative Neurology, 407(2), 183–192.

Mufson, E. J., & Pandya, D. N. (1984). Some observations on the course and composition of the cingulum bundle in the rhesus monkey. The Journal of Comparative Neurology, 225(1), 31– 43.

Neher, P., Hirjak, D., & Maier-Hein, K. (2023). Radiomic tractometry: a rich and tract-specific class of imaging biomarkers for neuroscience and medical applications. Research Square. 10.21203/rs.3.rs-2950610/v1

O’Donnell, L. J., Wells, W. M., 3rd, Golby, A. J., & Westin, C.-F. (2012). Unbiased groupwise registration of white matter tractography. Medical Image Computing and Computer-Assisted Intervention: MICCAI … International Conference on Medical Image Computing and Computer-Assisted Intervention, 15(Pt 3), 123–130.

O’Donnell, L. J., & Westin, C.-F. (2007). Automatic tractography segmentation using a high-dimensional white matter atlas. IEEE Transactions on Medical Imaging, 26(11), 1562–1575.

O’Donnell, L. J., Westin, C.-F., & Golby, A. J. (2009). Tract-based morphometry for white matter group analysis. NeuroImage, 45(3), 832–844.

O’Donnell, L., & Westin, C.-F. (2005). White matter tract clustering and correspondence in populations. Medical Image Computing and Computer-Assisted Intervention: MICCAI … International Conference on Medical Image Computing and Computer-Assisted Intervention, 8(Pt 1), 140–147.

Parente, P. M. D., & Santos Silva, J. M. (2016). Quantile Regression with Clustered Data. Journal of Econometric Methods, 5(1), 1–15.

Petrides, M., & Pandya, D. N. (2007). Efferent association pathways from the rostral prefrontal cortex in the macaque monkey. The Journal of Neuroscience: The Official Journal of the Society for Neuroscience, 27(43), 11573–11586.

Reddy, C. P., & Rathi, Y. (2016). Joint Multi-Fiber NODDI Parameter Estimation and Tractography Using the Unscented Information Filter. Frontiers in Neuroscience, 10, 166.

Reuben, D. B., Magasi, S., McCreath, H. E., Bohannon, R. W., Wang, Y.-C., Bubela, D. J., Rymer, W. Z., Beaumont, J., Rine, R. M., Lai, J.-S., & Gershon, R. C. (2013). Motor assessment using the NIH Toolbox. Neurology, 80(11 Suppl 3), S65–S75.

Ribeiro, M., Yordanova, Y. N., Noblet, V., Herbet, G., & Ricard, D. (2024). White matter tracts and executive functions: a review of causal and correlation evidence. Brain: A Journal of Neurology, 147(2), 352–371.

Richie-Halford, A., Yeatman, J. D., Simon, N., & Rokem, A. (2021). Multidimensional analysis and detection of informative features in human brain white matter. PLoS Computational Biology, 17(6), e1009136.

Ryan, M. C., Hong, L. E., Hatch, K. S., Gao, S., Chen, S., Haerian, K., Wang, J., Goldwaser, E. L., Du, X., Adhikari, B. M., Bruce, H., Hare, S., Kvarta, M. D., Jahanshad, N., Nichols, T. E., Thompson, P. M., & Kochunov, P. (2022). The additive impact of cardio-metabolic disorders and psychiatric illnesses on accelerated brain aging. Human Brain Mapping, 43(6), 1997–2010.

Sarica, A., Cerasa, A., Valentino, P., Yeatman, J., Trotta, M., Barone, S., Granata, A., Nisticò, R., Perrotta, P., Pucci, F., & Quattrone, A. (2017). The corticospinal tract profile in amyotrophic lateral sclerosis. Human Brain Mapping, 38(2), 727–739.

Schaeffer, D. J., Krafft, C. E., Schwarz, N. F., Chi, L., Rodrigue, A. L., Pierce, J. E., Allison, J. D., Yanasak, N. E., Liu, T., Davis, C. L., & McDowell, J. E. (2014). The relationship between uncinate fasciculus white matter integrity and verbal memory proficiency in children. Neuroreport, 25(12), 921–925.

Schilling, K. G., Archer, D., Yeh, F.-C., Rheault, F., Cai, L. Y., Hansen, C., Yang, Q., Ramdass, K., Shafer, A. T., Resnick, S. M., Pechman, K. R., Gifford, K. A., Hohman, T. J., Jefferson, A., Anderson, A. W., Kang, H., & Landman, B. A. (2022). Aging and white matter microstructure and macrostructure: a longitudinal multi-site diffusion MRI study of 1218 participants. Brain Structure & Function, 227(6), 2111–2125.

Schilling, K. G., Chad, J. A., Chamberland, M., Nozais, V., Rheault, F., Archer, D., Li, M., Gao, Y., Cai, L., Del’Acqua, F., Newton, A., Moyer, D., Gore, J. C., Lebel, C., & Landman, B. A. (2023). White matter tract microstructure, macrostructure, and associated cortical gray matter morphology across the lifespan. Imaging Neuroscience, 1, 1–24.

Schilling, K. G., Tax, C. M. W., Rheault, F., Landman, B. A., Anderson, A. W., Descoteaux, M., & Petit, L. (2022). Prevalence of white matter pathways coming into a single white matter voxel orientation: The bottleneck issue in tractography. Human Brain Mapping, 43(4), 1196–1213.

Schmahmann, J. D., & Pandya, D. (2009). Fiber pathways of the brain. Oxford University Press.

Smith, S. M., Jenkinson, M., Johansen-Berg, H., Rueckert, D., Nichols, T. E., Mackay, C. E., Watkins, K. E., Ciccarelli, O., Cader, M. Z., Matthews, P. M., & Behrens, T. E. J. (2006). Tract-based spatial statistics: voxelwise analysis of multi-subject diffusion data. NeuroImage, 31(4), 1487–1505.

Thiebaut de Schotten, M., Ffytche, D. H., Bizzi, A., Dell’Acqua, F., Allin, M., Walshe, M., Murray, R., Williams, S. C., Murphy, D. G. M., & Catani, M. (2011). Atlasing location, asymmetry and inter-subject variability of white matter tracts in the human brain with MR diffusion tractography. NeuroImage, 54(1), 49–59.

Tulsky, D. S., Carlozzi, N., Chiaravalloti, N. D., Beaumont, J. L., Kisala, P. A., Mungas, D., Conway, K., & Gershon, R. (2014). NIH Toolbox Cognition Battery (NIHTB-CB): list sorting test to measure working memory. Journal of the International Neuropsychological Society: JINS, 20(6), 599–610.

Van Essen, D. C., Smith, S. M., Barch, D. M., Behrens, T. E. J., Yacoub, E., Ugurbil, K., & WU-Minn HCP Consortium. (2013). The WU-Minn Human Connectome Project: an overview. NeuroImage, 80, 62–79.

Vázquez, A., López-López, N., Sánchez, A., Houenou, J., Poupon, C., Mangin, J.-F., Hernández, C., & Guevara, P. (2020). FFClust: Fast fiber clustering for large tractography datasets for a detailed study of brain connectivity. NeuroImage, 220, 117070.

Vos, S. B., Viergever, M. A., & Leemans, A. (2013). Multi-fiber tractography visualizations for diffusion MRI data. PloS One, 8(11), e81453.

Wasserthal, J., Neher, P. F., Hirjak, D., & Maier-Hein, K. H. (2019). Combined tract segmentation and orientation mapping for bundle-specific tractography. Medical Image Analysis, 58(101559), 101559.

Wasserthal, J., Neher, P., & Maier-Hein, K. H. (2018). TractSeg - Fast and accurate white matter tract segmentation. In arXiv [cs.CV] (pp. 239–253). arXiv. 10.1016/j.neuroimage.2018.07.070

Wei, Y., Pere, A., Koenker, R., & He, X. (2006). Quantile regression methods for reference growth charts. Statistics in Medicine, 25(8), 1369–1382.

Welniarz, Q., Dusart, I., & Roze, E. (2017). The corticospinal tract: Evolution, development, and human disorders. Developmental Neurobiology, 77(7), 810–829.

Yakovlev, P. I., & Locke, S. (1961). Limbic nuclei of thalamus and connections of limbic cortex. III. Corticocortical connections of the anterior cingulate gyrus, the cingulum, and the subcallosal bundle in monkey. Archives of Neurology, 5(4), 364–400.

Yeatman, J. D., Dougherty, R. F., Myall, N. J., Wandell, B. A., & Feldman, H. M. (2012). Tract profiles of white matter properties: automating fiber-tract quantification. PloS One, 7(11), e49790.

Yeatman, J. D., Dougherty, R. F., Rykhlevskaia, E., Sherbondy, A. J., Deutsch, G. K., Wandell, B. A., & Ben-Shachar, M. (2011). Anatomical properties of the arcuate fasciculus predict phonological and reading skills in children. Journal of Cognitive Neuroscience, 23(11), 3304–3317.

Yeh, F.-C. (2020). Shape analysis of the human association pathways. NeuroImage, 223, 117329.

Yushkevich, P. A., Zhang, H., Simon, T. J., & Gee, J. C. (2009). Structure-specific statistical mapping of white matter tracts. In Mathematics and Visualization (pp. 83–112). Springer Berlin Heidelberg.

Zekelman, L. R., Zhang, F., Makris, N., He, J., Chen, Y., Xue, T., Liera, D., Drane, D. L., Rathi, Y., Golby, A. J., & O’Donnell, L. J. (2022). White matter association tracts underlying language and theory of mind: An investigation of 809 brains from the Human Connectome Project. NeuroImage, 246, 118739.

Zelazo, P. D., Anderson, J. E., Richler, J., Wallner-Allen, K., Beaumont, J. L., Conway, K. P., Gershon, R., & Weintraub, S. (2014). NIH Toolbox Cognition Battery (CB): validation of executive function measures in adults. Journal of the International Neuropsychological Society: JINS, 20(6), 620–629.

Zhang, F., Daducci, A., He, Y., Schiavi, S., Seguin, C., Smith, R. E., Yeh, C.-H., Zhao, T., & O’Donnell, L. J. (2022). Quantitative mapping of the brain’s structural connectivity using diffusion MRI tractography: A review. NeuroImage, 249, 118870.

Zhang, F., Wu, Y., Norton, I., Rathi, Y., Golby, A. J., & O’Donnell, L. J. (2019). Test-retest reproducibility of white matter parcellation using diffusion MRI tractography fiber clustering. Human Brain Mapping, 40(10), 3041–3057.

Zhang, F., Wu, Y., Norton, I., Rigolo, L., Rathi, Y., Makris, N., & O’Donnell, L. J. (2018). An anatomically curated fiber clustering white matter atlas for consistent white matter tract parcellation across the lifespan. NeuroImage, 1(179), 429–447.

Zhang, H., Schneider, T., Wheeler-Kingshott, C. A., & Alexander, D. C. (2012). NODDI: practical in vivo neurite orientation dispersion and density imaging of the human brain. NeuroImage, 61(4), 1000–1016.

